# Molecular mechanism of plasmid elimination by the DdmDE defense system

**DOI:** 10.1101/2024.05.10.593530

**Authors:** L. Loeff, D.W. Adams, C. Chanez, S. Stutzmann, L. Righi, M. Blokesch, M. Jinek

**Author notes:** **Correspondence:** &.

## Abstract

Seventh pandemic *Vibrio cholerae* strains contain two hallmark pathogenicity islands that encode the DNA defense modules DdmABC and DdmDE. Here we use cryo-EM to reveal the mechanistic basis for plasmid defense by DdmDE. A cryo-EM structure of the DdmD helicase-nuclease reveals that it adopts an auto-inhibited dimeric architecture. The prokaryotic Argonaute protein DdmE uses a DNA guide to target plasmid DNA. A structure of the DdmDE complex, validated by *in vivo* mutational studies, shows that DNA binding by DdmE triggers disassembly of the DdmD dimer and loading of monomeric DdmD onto the non-target DNA strand. Finally, *in vitro* studies reveal that DdmD translocates in the 5’ to 3’ direction, while partially degrading the plasmid DNA. These findings provide critical insights into the architecture and mechanism of DdmDE systems in plasmid elimination.

## Introduction

The evolutionary arms race between prokaryotes and invasive mobile genetic elements (MGEs) has resulted in the emergence of a myriad of host-defense systems that provide immunity against invading MGEs (*1*). These immune mechanisms include restriction-modification (R-M), CRISPR-Cas, Argonaute, CBASS, Shedu, Lamassu and Wadjet systems (*2–10*). Defense systems play a key role in the elimination of invading MGE and shape microbial communities and ecosystems by limiting horizontal gene transfer (HGT) (*11, 12*). As numerous molecular genetic engineering tools have their origins in prokaryotic genome defense systems, understanding prokaryotic immune systems is not only crucial for unraveling the dynamics of prokaryotic host-MGE interactions but also for the development of molecular tools with applications in biotechnology and medicine.

In the important human pathogen *Vibrio cholerae*, two DNA defense modules termed DdmABC and DdmDE cooperate to eliminate plasmids and are thought to have played a key role in the evolutionary success of the seventh pandemic O1 El Tor (7PET) strains (*13*). DdmABC is a Lamassu-like defense system, which has been shown to trigger abortive infection upon activation by plasmids and bacteriophages (*7, 13, 14*). By contrast, the DdmDE system acts directly against small plasmids, resulting in their degradation (*13*). Structural modelling suggests DdmE is a prokaryotic Argonaute (pAgo) protein that might recognize plasmid DNA, while DdmD is predicted to encode a fusion protein comprising helicase and nuclease domains that could mediate plasmid degradation (*13*). Although the functional role of the DdmDE system in plasmid elimination has been established, the mechanistic basis for its function remains elusive.

## Results

### DdmE is DNA-guided DNA-targeting Argonaute

pAgos use short nucleic acid guides to direct the recognition and targeting of nucleic acids. To obtain mechanistic insights into the function of the pAgo protein DdmE, we initially performed a phylogenetic analysis of 828 pAgo sequences, comprising DdmE orthologs and both short and long pAgo proteins (*9, 23, 24*). DdmE orthologs clustered with the long pAgos, forming a sub-clade that is distinct from both long-A and long-B pAgos (**Fig. 1A**), hereafter referred to as long-C. Multiple sequence alignments and AlphaFold 2 structural modeling of DdmE revealed the absence of a canonical DEDX catalytic tetrad in its P-element-induced wimpy testis (PIWI) domain (**Fig S1A-B**), indicating that DdmE proteins lack endonuclease activity and suggesting that the DdmE does not rely on endonuclease-catalyzed target nucleic acid cleavage, in contrast to long-A pAgo proteins.

**Figure 1:**
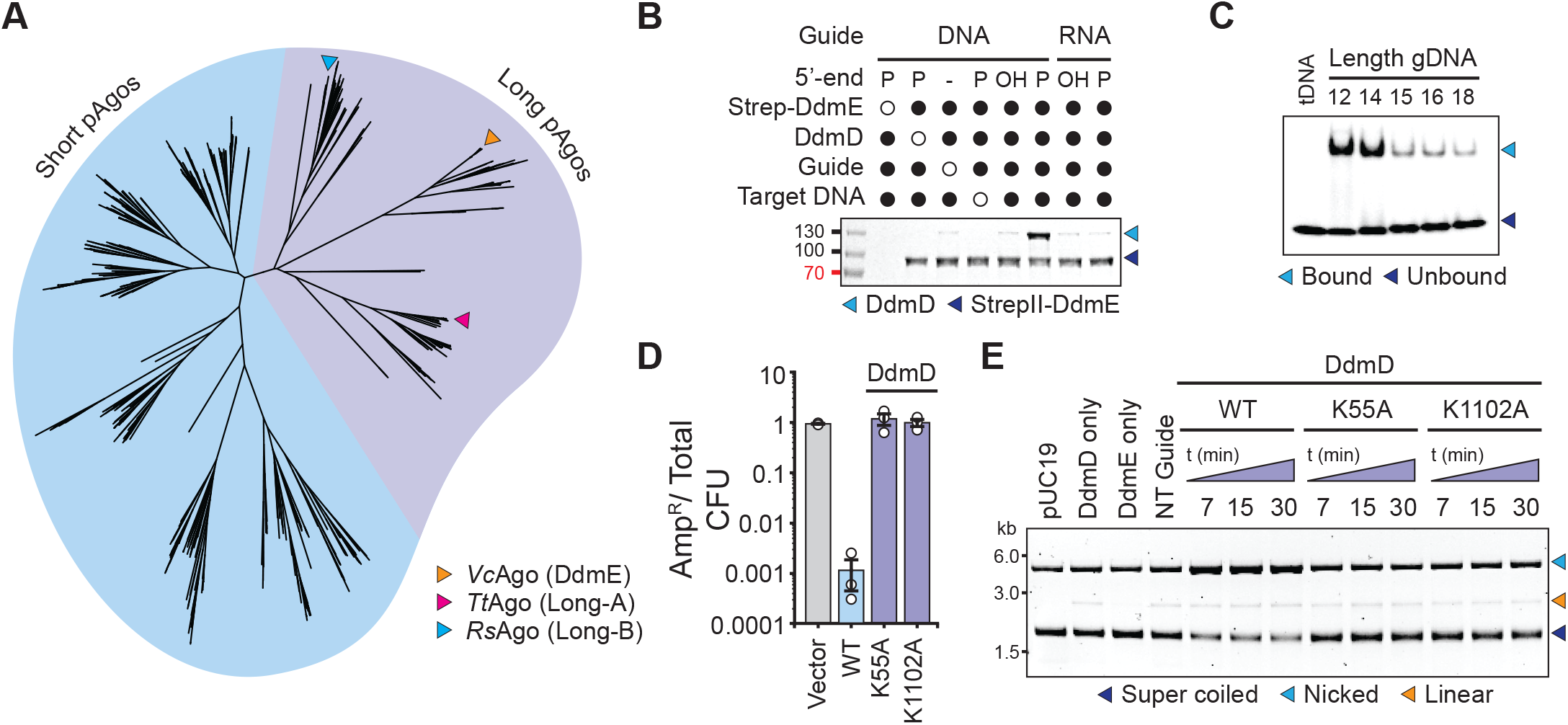
DdmE pAgo uses DNA guides to recruit DdmD helicase-nuclease. (**A**) Phylogenetic classification of 828 prokaryotic Argonautes that cluster into short and long pAgos. DdmE orthologs cluster in a distinct clade that are different from Long-A and Long-B pAgos. (**B**) *In vitro* pulldown experiment with recombinant DdmD and StrepII-tagged DdmE in the presence of 14-nt guides carrying 5’-phosphate (P) or 5’-hydroxyl (OH) groups and a dsDNA target containing a mismatched region over the target site (bubble). (**C**) Electrophoretic mobility shift analysis of DdmE binding to fluorescently labelled ssDNA target in the presence of with 5’-phopshorylated DNA guides (gDNAs) of different length. (**D**) *In vivo* plasmid elimination assay in *E. coli* expressing wild-type (WT) DdmDE or WT DdmE and DdmD mutants, or a vector-only control. Data represents mean fraction of plasmid-bearing colony forming units (CFU) ± SEM of three independent replicates (n=3). (**E**) *In vitro* plasmid degradation assays with WT DdmDE or DdmD proteins carrying inactivating mutations in the helicase (K55A) or nuclease (K1102A) domains. DdmD-only, DdmE-only and non-targeting guide (NT guide) controls were incubated for the entire duration of the experiment. ± SEM of three biological replicates (n=3).

Size exclusion chromatography analysis of purified recombinant *V. cholerae* DdmD and DdmE proteins revealed that DdmE is a monomer in solution, while DdmD forms a homodimer (**Fig. S2A-C**). As we were unable to detect a stable interaction between DdmD and DdmE in isolation (**Fig. S2D**), we hypothesized that DdmE facilitates the recruitment of DdmD to a D-loop structure generated by guide-dependent target nucleic acid binding. To test this, we performed affinity co-precipitation experiments of DdmD using matrix-immobilized DdmE in the presence of DNA or RNA guides and targets. When DdmE was loaded with a 14-nucleotide (nt), 5′-phosphorylated single-stranded (ss) DNA guide and incubated with a double-stranded (ds) DNA target containing internal mismatched sequences on the non-targeted DNA strand within the target site (to facilitate formation of a D-loop structure), we observed efficient DdmD co-precipitation. DdmD was not co-precipitated by DdmE in the presence a guide DNA lacking a 5’-phosphate group or with RNA guides (**Fig 1B**). Moreover, DdmD co-precipitation did not occur when the target was ssRNA, ssDNA or a perfectly paired dsDNA (**Fig S2E-G**). These results suggest that the DdmDE system uses DdmE loaded with 5’-phosphorylated DNA guides to target DNA and recruit DdmD.

pAgo proteins have been shown to use guides of varying length. Using an electrophoretic mobility shift assay with a fluorophore-labeled target ssDNA and 5′-phosphorylated guide DNAs, we determined that the optimal length of DdmE guides is in the range of 12-14 nt (**Fig. 1C**). Notably, no cleavage of the target DNA was observed in this assay (**Fig. S2H**), indicating that DdmE is catalytically inactive in agreement with the absence of the canonical DEDX catalytic tetrad in its PIWI domain (**Fig S1A-B**). Altogether, these data indicate that DdmE uses short (<15 nt) 5’-phosphorylated DNAs as guides for target DNA binding.

### Plasmid elimination requires ATPase and nuclease activities of DdmD

Structural modeling predicted that DdmD contains an N-terminal super family 2 (SF2) helicase and C-terminal PD-(D/E)xK nuclease domain (*13*), suggesting that DdmD may act as a downstream effector that degrades plasmid DNA upon recruitment and activation by DdmE. To test this hypothesis, we utilized an *in vivo* assay that probes target plasmid elimination in *Escherichia coli* after growth for 10 generations (**Fig S2I**). Induction of DdmDE expression resulted in an approximately three-order-of-magnitude decrease of number of plasmid-bearing colonies as compared to uninduced and vector-only controls (**Fig. 1D** and **Fig S2I**). In contrast, mutation of the helicase (K55A, designed to impair ATP binding) or nuclease domains (K1102A) of DdmD abrogated plasmid elimination by the DdmDE system (**Fig. 1D**), consistent with our biochemical analysis and our previously reported mutations of *ddmD* at its native genomic locus in *V. cholerae* (*13*). These results confirm that both helicase and nuclease activity of DdmD are essential for plasmid elimination by the DdmDE system.

To validate these observations biochemically, we reconstituted plasmid elimination by the DdmDE system *in vitro*. Upon incubation of a target plasmid with DdmD and DdmE proteins in the presence of a complementary guide DNA and ATP, the supercoiled plasmid DNA was rapidly converted to nicked cleavage products (**Fig 1E**). Plasmid nicking and cleavage was not observed with DdmD proteins containing inactivating mutations in the ATPase (K55A, **Fig. 1E** and **S2J**) or nuclease domains (K1102A, **Fig. 1E** and **S2K**). Notably, DdmD alone showed substantial degradation activity on ssDNA plasmids (**Fig. S2K)**, while no degradation of the plasmid dsDNA was observed with DdmD alone (**Fig. 1E**), indicating that DdmD is a potent ssDNA nuclease that requires DdmE for activation and recruitment to its plasmid DNA target. Together, these results indicate that the activity of the DdmDE system involves DNA-guided targeting by DdmE and ATPase-dependent, nuclease-catalyzed plasmid degradation by DdmD.

### DdmD is autoinhibited by dimerization

To gain mechanistic insights into the activity of the helicase-nuclease DdmD dimer within the DdmDE system, we determined the atomic structure of dimeric DdmD by single-particle cryo-EM, obtaining a reconstruction with a nominal resolution of 2.6 Å (**Fig. 2A** and **S3A-D**). The structure reveals a C2 homodimeric assembly in which the individual protomers dimerize via an extensive electrostatic interface that spans a surface area of ∼2498 Å^2^ (**Fig. 2B**). The domain organization of the DdmD N-terminal helicase closely resembles that of *E. coli* DinG, a processive 5’-3’ translocating SF2 DNA helicase that belongs to the XPD family (**Fig. 2C** and **Fig. S4A**) (*15, 16*). The helicase core of DdmD is composed of two canonical RecA-like helicase domains (HD1 and HD2), which together form an ATPase active site located at their interface (**Fig. 2C**). The HD1 domain further houses two inserted domains, hereafter called INS and Arch, that resemble the iron sulfur cluster and arch domains found in XPD type helicases, respectively (**Fig. 2C** and **Fig. S4B-C**). Unlike DinG, the DdmD INS domain lacks an iron-sulfur cluster.

**Figure 2:**
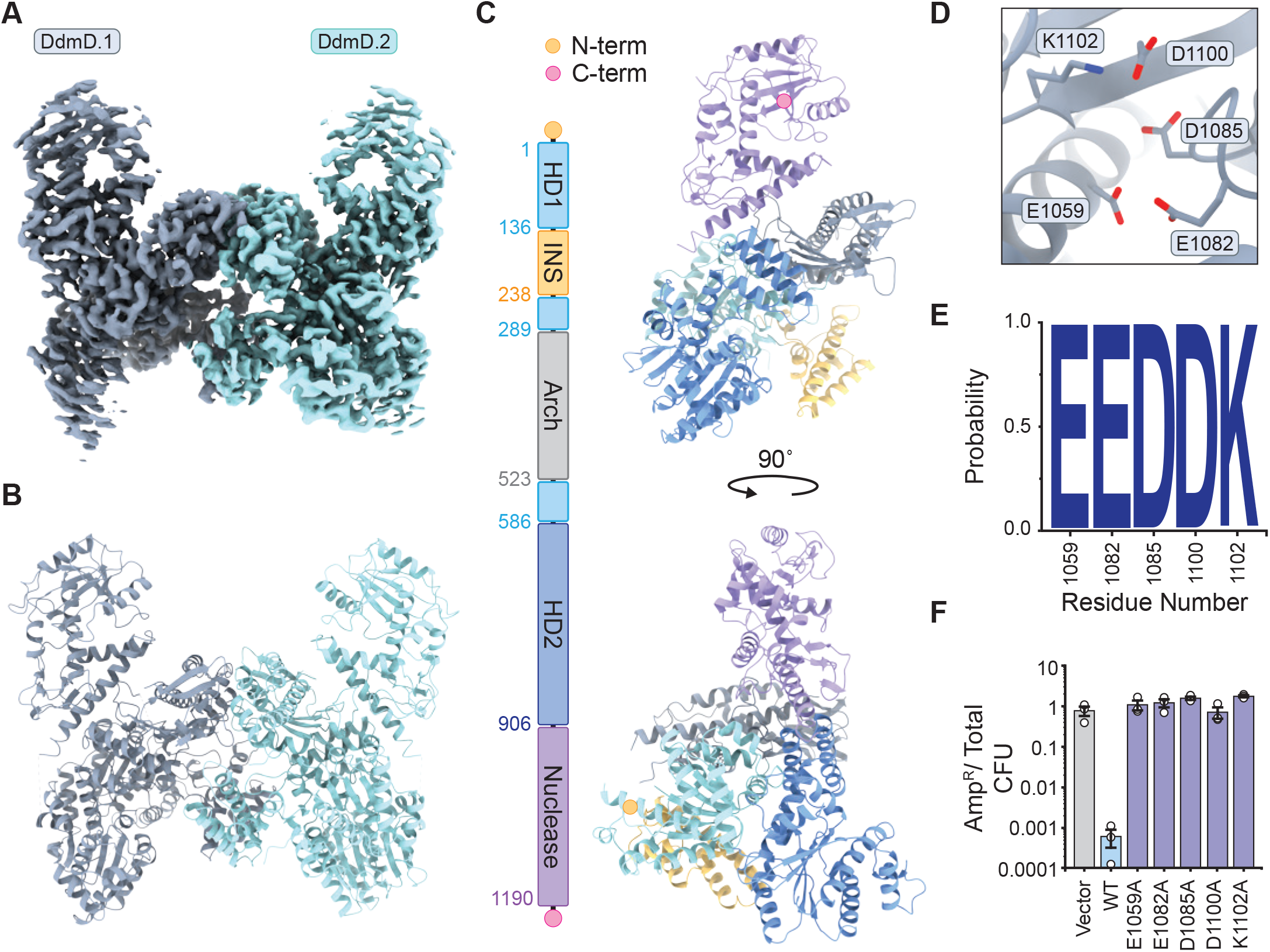
DdmD forms an autoinhibited dimer. (**A**) Cryo-EM density map of the dimeric DdmD complex. (**B**) Structural model of tetrameric DdmD, shown in cartoon representation in the same orientation as in A. (**C**) Structural model of a DdmD protomer (right) shown alongside domain organisation (left). N- and C-termini are indicated by orange and pink circles, respectively. (**D**) Close-up view of the PD-(D/E)xK nuclease active site of DdmD showing its catalytic residues. (**E**) Sequence conservation analysis of nuclease active site residues in DdmD. Figure was generated using WebLogo (*46*). (**F**) *In vivo* plasmid elimination assay in *E. coli* strains expressing WT DdmDE, WT DdmE and DdmD active site mutants, or a vector-only control. Data represents mean fraction of plasmid-bearing colony forming units (CFU) ± SEM of three independent replicates (n=3). Experimental data was acquired simultaneously with data displayed in Fig S7D and share the same WT and vector-only controls.

The INS and Arch domains form the dimerization interface within the DdmD, and together with the HD1 and HD2 domains surround a central DNA binding channel (**Fig. 2B** and **S4A**). Superposition of the DdmD dimer with the structure of DinG in complex with ssDNA and an ATP mimic (PDB: 6FWS) reveals that the dimer interface occludes the 3’ end of the DNA binding channel, thereby precluding ATP-driven translocation along the DNA (**Fig S4D**). This suggests that DdmD exists in an autoinhibited state in its dimeric form, consistent with the observation that DdmD does not exhibit nuclease activity in vitro in the absence of DNA-guided DdmE. The C-terminal domain of DdmD is positioned above the helicase core and resembles the signature fold of enzymes belonging to the PD-(D/E)xK phosphodiesterase superfamily (**Fig. 2C** and **S4E**) (*17*), containing a conserved catalytic pocket lined with residues Glu1059^DdmD^, Glu1082^DdmD^, Asp1085^DdmD^, Asp1100^DdmD^ and Lys1102^DdmD^ (**Fig 2D-E** and **S4F**). Alanine substitutions of these residues abolished DdmDE-dependent plasmid elimination in both *E. coli* and in *V. cholerae* (**Fig. 2F** and **S4G**), confirming that DdmD is a PD-(D/E)xK-like nuclease whose catalytic activity is essential for plasmid elimination by the DdmDE defense system.

### DNA-guided target DNA recognition and DdmD recruitment by DdmE

To obtain structural insights into the molecular architecture of DdmE and its role in the activation of DdmD, we reconstituted a DdmD-DdmE complex with a 14-nt 5′-P DNA guide and a target dsDNA with a mismatched non-target strand (NTS) to facilitate D-loop formation (**Fig. 1B** and **3A**). Subsequent single-particle cryo-EM analysis resulted in a reconstruction with a nominal resolution of 2.6 Å (**Fig. 3B** and **S5A-D**). The structure of the DdmDE complex reveals that the DdmD homodimer disassembles upon interaction with target-bound DdmE, forming a 1:1 heterodimer in which DdmE is bound to the guide DNA (gDNA)-target DNA strand (tDNA) duplex, while DdmD engages the displaced non-target DNA strand (ntDNA) (**Fig. 3B-C**).

**Figure 3:**
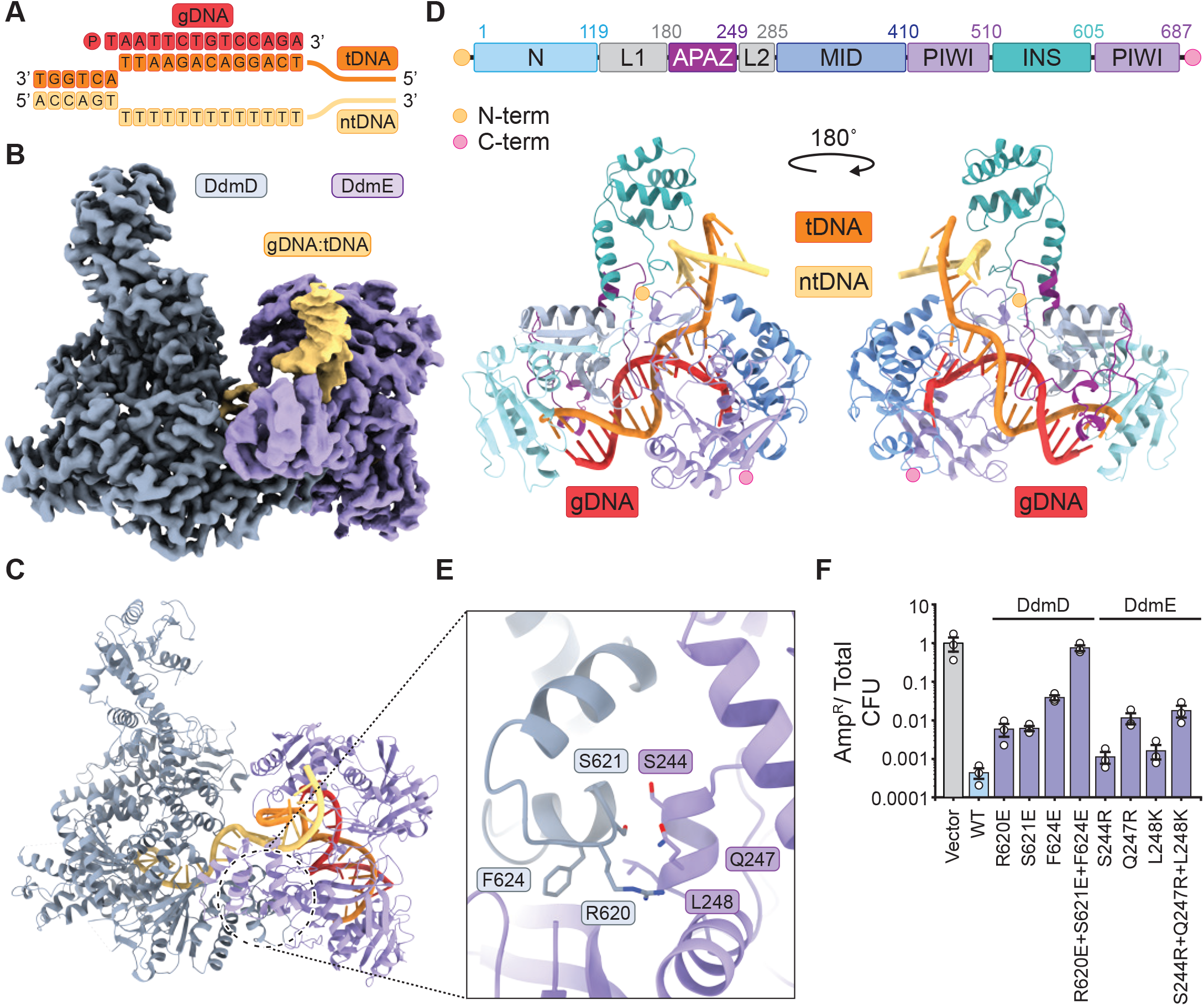
Molecular basis of DdmD recruitment by DdmE. (**A**) Schematic diagram of the dsDNA bubble substrate used for the cryo-EM complex assembly. Nucleotides that are visible in the cryo-EM structure are displayed as letters. The 14-nt guide DNA (gDNA), target DNA strand (tDNA) and non-targeted strand (ntDNA) are highlighted in red, orange, and yellow, respectively. (**B**) Cryo-EM density map of the DdmD-DdmE holocomplex. (**C**) Structural model of the DdmD-DdmE holocomplex. (**D**) Structural model of the DdmE monomer within the DdmD-DdmE holocomplex (bottom) shown alongside domain organisation (top). Part of the non-target DNA strand has been removed for clarity. (**E**) Close-up view of the DdmD-DdmE interaction that is centered on an alpha-helical bundle in the DdmD HD domain. (**F**) *In vivo* plasmid elimination assay in *E. coli* strains expressing WT DdmDE system, DdmD and DdmE interface mutants, or a vector-only control. Data represents mean fraction of plasmid-bearing colony forming units (CFU) ± SEM of three independent replicates (n=3).

DdmE adopts a conserved domain architecture observed in other pAgo proteins, comprising the N-terminal, L1, L2, MID, PIWI domains, except that it lacks a canonical PAZ domain and instead contains a smaller domain, conserved in members of the long-C pAgo clade, herein referred to as C-APAZ (long-C analog of PAZ) (**Fig. 3D**). Additionally, the PIWI domain contains a domain insertion (INS), also conserved within the long-C pAgos (**Fig. S6A-B**). The 5′-terminal nucleotide of the guide DNA is unpaired and sequestered in a highly conserved pocket within the MID domain by interactions with Tyr363^DdmE^ and Lys405^DdmE^ (**Fig. S7A-C**). The 5’-terminal and third backbone phosphate groups of the guide DNA are coordinated by a Mg^2+^ ion, in turn contacted by residues Gln376 ^DdmE^, Glu401^DdmE^, Lys674^DdmE^ and Glu678^DdmE^ (**Fig. S7A-B**). In agreement with the observed mode of guide DNA binding, alanine substitutions of the above residues resulted in a complete loss of plasmid elimination in *E. coli* (**Fig. S7D**). In *V. cholerae* harboring both the DdmABC and DdmDE systems, these mutations resulted in complete or partial loss of plasmid elimination, while complete loss of plasmid elimination was observed for all DdmE mutants in a strain lacking the cooperating system DdmABC (**Fig. S7E-F**).

The distal end of the gDNA-tDNA duplex is capped by the N domain of DdmE. Structural modeling suggests that extension of the guide beyond 14-bp would result in a steric clash with the N domain (**Fig. S8A**), explaining the strong decrease in target binding observed for guide DNAs longer than 14-nt (**Fig. 1C)**. DdmE interacts with the tDNA at the proximal D-loop junction with the gDNA by salt-bridge interactions with the tDNA phosphate backbone (via Lys625^DdmE^ and Arg664^DdmE^) and hydrogen bonding and π-π stacking contacts with tDNA nucleobases (via His393^DdmE^, Arg662^DdmE^ and His663^DdmE^), all of which are required for efficient plasmid elimination by the DdmDE system in *E. coli* (**Fig. S8B-D**). The C-APAZ domain interrogates the minor groove the gDNA-tDNA duplex using Lys230^DdmE^ and Arg232^DdmE^ and simultaneously interacts with DdmD (**Fig. S8E)**. This suggests that the C-APAZ domain acts as a sensor that recognizes B-form duplex geometry upon guide and target DNA hybridisation to mediate DdmD recruitment.

DdmD uses its HD2 domain to interact with DdmE through an electrostatic interface that spans a surface area of ∼1160 Å^2^ (**Fig. 3C** and **S9A**). The interaction involves an alpha-helical bundle (residues Ser603^−^Ala633^DdmD^) within HD2^DdmD^ that is wedged between the INS and C-APAZ domains of DdmE (**Fig. 3E** and **S9B**). Consistent with these observations, mutations of the interacting residues in DdmD and DdmE resulted in moderate to strong reduction of anti-plasmid activity in *E. coli* (**Fig. 3F** and **S9E**). The DdmDE interaction is further reinforced by electrostatic interactions between a basic patch on DdmD and an acidic patch on the INS domain of DdmE (**Fig. S9A-D)**, as confirmed by the loss of anti-plasmid activity upon introduction of charge-neutralizing mutations in the DdmD basic patch, or deletion of the DdmE INS domain (**Fig. S9E)**. Collectively, these findings show that target DNA-bound DdmE recruits DdmD and reveals critical molecular determinants of the DdmDE interaction necessary to support the anti-plasmid activity of the system.

### DdmE mediates DdmD loading onto non-target DNA strand

The structure of the DdmD-DdmE complex reveals a partial D-loop in which DdmE is bound to the tDNA strand in a guide-dependent manner, while DdmD engages the displaced ntDNA strand (**Fig. 3A** and **Fig. 4A**). Starting from the flanking duplex part of the D-loop, the first two ntDNA nucleotides (dT7–8) are contacted by DdmE. The ntDNA strand is then kinked at dT9 as it enters the HD2 domain of DdmD, with the nucleobases of dT9–dT11 forming a continuous stack (**Fig. 4A-B**). At nucleotide dT11, the ntDNA is kinked again as it makes a π-π stacking interaction with Phe639^DdmD^ and passes through a constriction formed by Gln781^DdmD^ and Arg828^DdmD^. The downstream nucleotides form two continuous base-stacked segments as the ntDNA spans from HD2 (dT12-dT15) to HD1 (dT16-dT19). The 3’-terminal nucleotide (dT19) is stabilized by Tyr194^DdmD^ via π-π stacking as the ntDNA strand exits DdmD (**Fig. 4C**). In agreement with the structural observations, alanine substitutions of the residues at the proximal end of the D-loop resulted in a complete loss of plasmid elimination in *E. coli*, while substitution of Tyr194^DdmD^ resulted in a minor reduction of anti-plasmid activity (**Fig. 4D**).

**Figure 4:**
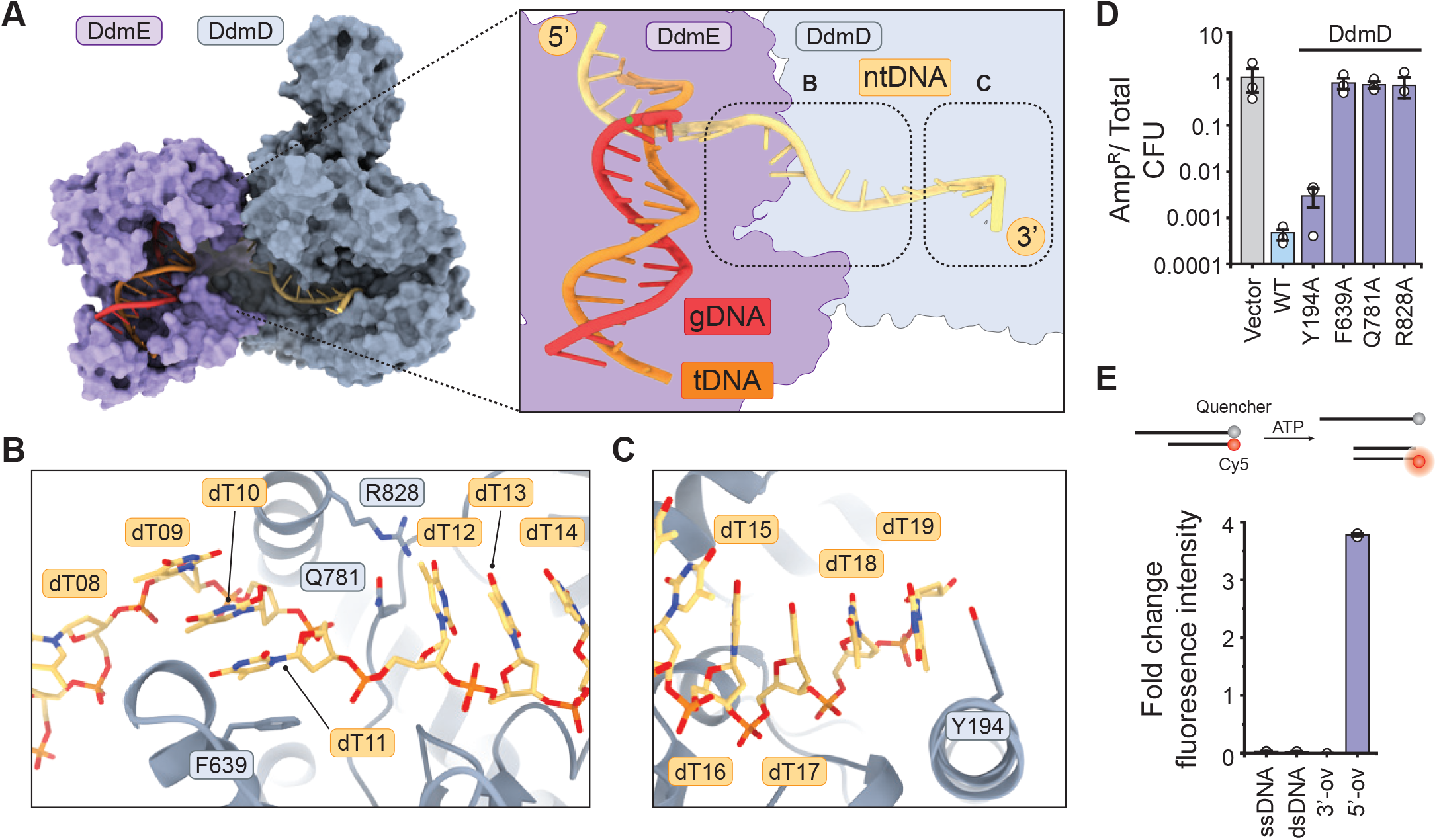
DdmD is loaded on non-target DNA strand for 5’-3’ translocation. (**A**) Surface representation of the DdmD-DdmE holo complex (right). Inset shows zoomed-in view of the structurally ordered part of the D-loop comprising guide DNA (gDNA)-target DNA strand (tDNA) duplex and the non-target DNA. (**B**) Close-up view of DdmD interactions with the proximal end of the non-target DNA strand. (**C**) Close-up view of DdmD interactions with the distal end of the non-target DNA strand. (**D**) *In vivo* plasmid elimination assay in *E. coli* strains expressing WT DdmDE system, WT DdmE and DdmD mutants, or a vector-control. Data represents mean plasmid bearing colony forming units (CFU) ± SEM of three independent replicates (n=3). Experimental data was acquired simultaneously with data displayed in Fig S9E and share the same WT and vector only controls. (**E**) Top: Schematic showing a fluorophore-quencher DNA unwinding assay to probe the helicase polarity of DdmD using ssDNA, or dsDNA substrates with blunt ends, a 5’ overhang (5’-ov) or a 3’ overhang (3’-ov). Bottom: Helicase activity of DdmD, quantified by change in fluorescence intensity upon incubation of the DNA substrate with DdmD and ATP.

Although not directly observed in the cryo-EM map, the ntDNA presumably continues beyond the exit site, thereby likely promoting disruption of the DdmD dimer upon interaction with target-bound DdmE. Furthermore, the observed polarity of the bound ntDNA is consistent with the unwinding mechanism of XDP-like helicases (*16*), suggesting that DdmD is a translocating DNA helicase with a 5’ to 3’ directionality. To verify this, we performed DNA unwinding assays using fluorophore-quencher DNA substrates (**Fig 4E**). Unwinding activity was only observed with a duplexed DNA containing a 5’-overhang, confirming that DdmD is a 5’-3’ unwinding helicase (**Fig 4E**). Together, these structural and biochemical findings indicate that upon tDNA binding, DdmE recruits DdmD to the non-target DNA strand, which results in DdmD activation and 5’-3’ translocation along the ntDNA.

## Discussion

Here we shed light on the molecular mechanisms underpinning the anti-plasmid activity of the DdmDE system of *V. cholerae*. We show that DdmE, which belongs to a novel clade of catalytically inactive long pAgos, employs 5’-phosphorylated DNA guides to target DNA. Target binding by DdmE recruits the helicase-nuclease protein DdmD, which initially resides in an auto-inhibited state due to self-dimerization. Our findings suggest a mechanistic model (**Fig. 5**), in which the DdmD dimer disassembles upon loading of one protomer onto the non-target DNA strand. This in turn primes DdmD for degradation of the non-target DNA by ATP-driven, processive translocation in a 5’ to 3’ direction. We speculate that DdmD and DdmE remain in complex upon translocation, generating ssDNA loops that can efficiently be cleaved by the PD-(D/E)xK nuclease domain of DdmD to generate small ssDNA gaps or nicks in the target plasmid. In this way, DdmDE is able to interfere with plasmid replication and possibly target the plasmid DNA for degradation by host nucleases. We cannot exclude the possibility that DdmD degrades the non-target DNA strand exonucleolytically after initial nicking, although this was not observed *in vitro*. DdmE appears to lack an intrinsic ability to unwind dsDNA targets, as for other DNA targeting pAgo proteins (*18*), possibly relying on negative supercoiling of plasmid DNA *in vivo*.

**Figure 5:**
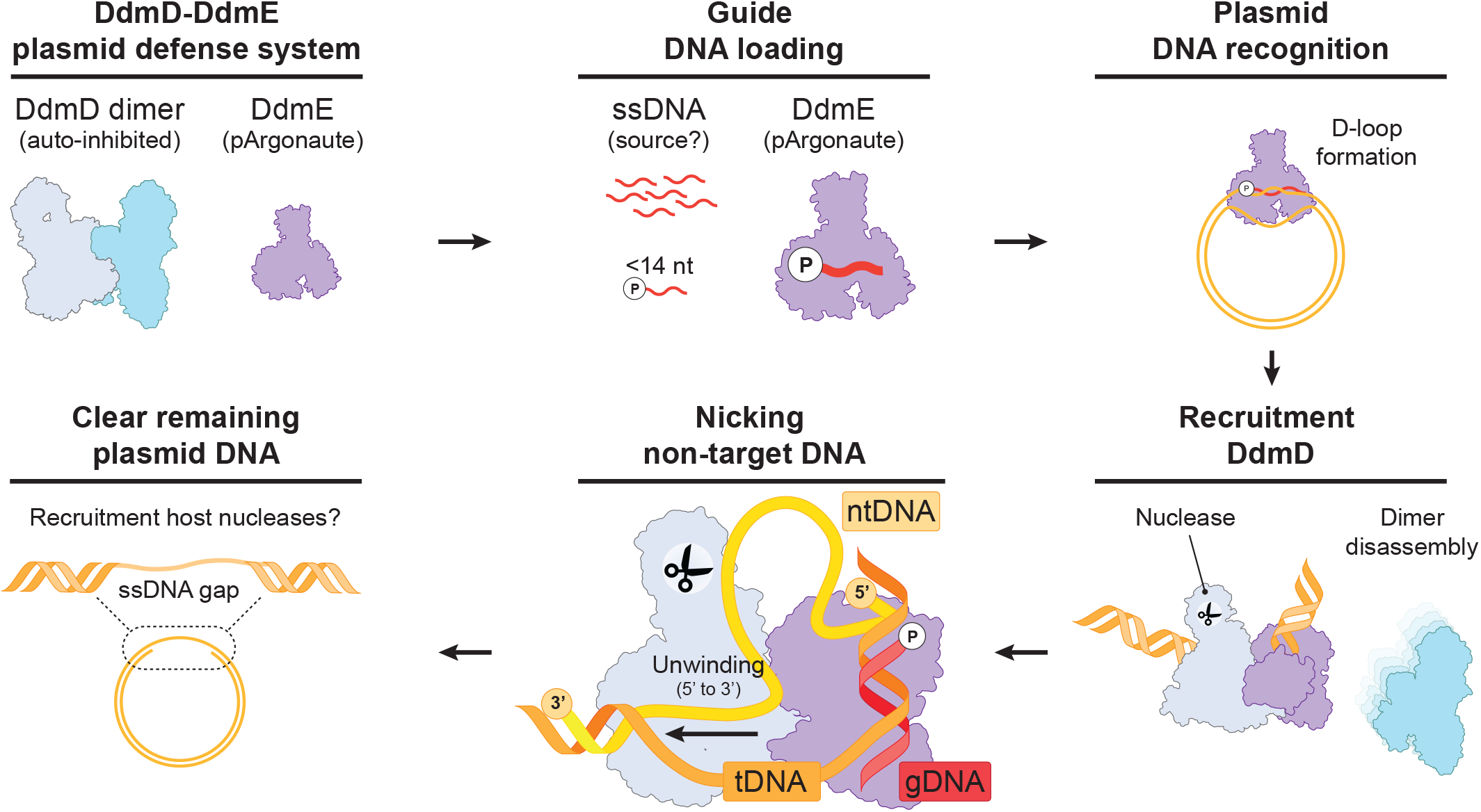
Mechanistic model for plasmid elimination by DdmDE. DdmE is guided by short DNAs with a 5’ terminal phosphate to bind plasmid DNA targets. Upon target DNA binding and D-loop formation, DdmE recruits DdmD and facilitates its loading onto non-target DNA strand. The autoinhibited DdmD dimer disassembles upon recruitment to target bound DdmE. DdmD translocates in a 5’ to 3’ direction on the non-targeted strand, feeding the single-stranded non-target strand into the nuclease domain of DdmD. This generates nicks or ssDNA gaps that interfere with plasmid propagation or facilitate further degradation by host nucleases.

The mechanism of DdmDE brings up notable parallels with the molecular mechanism of type I CRISPR-Cas systems, in which the RNA-guided effector complex Cascade recruits the helicase-nuclease fusion protein Cas3 to the non-target DNA strand (*19, 20*). Subsequently, Cascade and Cas3 remain in complex while the DNA is repeatedly unwound by the helicase domain of Cas3, generating short stretches of ssDNA in the DNA target (*21, 22*). The origin of the guide DNAs used by DdmDE remains an outstanding question and we hypothesize that guide generation is coupled to DNA end resection by the double-strand break repair complex RecBCD (*23*), which has been shown to be a source of guides for other pAgos and CRISPR-Cas systems (*24, 25*). The dependence on RecBCD could further explain the bias in immunity for small multicopy plasmids that lack Chi sites as compared to chromosomal DNA.

Our work demonstrates that DdmE mutant variants that are non-functional for plasmid elimination in *E. coli* remain active in a *V. cholerae* strain harboring the Lamassu-like DdmABC system. While understanding the functional interactions between DdmABC and DdmDE will require deeper exploration of the molecular mechanisms of DdmABC and its cooperation with DdmDE, these findings further reinforce the functional link between these systems in their native host. The collaboration between DdmABC and DdmDE aligns with studies showing that co-occurring bacterial defense systems synergize to provide anti-phage activities (*26*), emphasizing the importance of this concept in prokaryotic immunity. Furthermore, defining the cooperation of DdmABC and DdmDE in anti-plasmid defense is important for understanding the epidemiological success of the seventh-pandemic *V. cholerae* strains.

Given that type I CRISPR-Cas systems have been repurposed as molecular targeting technologies (*27, 28*), our study suggests that DdmDE may be exploited to deplete bacterial strains though the delivery of DNAs that would act as sources of self-targeting guides. A similar approach based on CRISPR-Cas9 has been used to target virulence genes in pathogenic *Staphylococcus aureus* strains (*29*). Alternatively, DdmDE might be utilized to generate self-eliminating plasmids for recombi-neering-based genome engineering approaches. Taken together, our study provides fundamental mechanistic insights into the function of DdmDE systems in genome defense and might pave the way towards harnessing them as novel antimicrobial or genetic engineering tools.

## Acknowledgements

We are grateful to Piotr Szwedziak (University of Zurich Center for Microscopy and Image Analysis) for technical support with cryo-EM data acquisition. We thank all the members of the Jinek and Blokesch labs for constructive feedback throughout the project.

## Author Contributions

Conceptualization: LL, DWA, MB, MJ

Methodology: LL, DWA

Investigation: LL, DWA, CC, SS, LR, MB

Visualization: LL

Funding acquisition: MB, MJ

Supervision: LL, MB, MJ

Writing – original draft: LL

Writing – review & editing: LL, DWA, MB, MJ

## Data and materials availability

Atomic coordinates and cryo-EM maps for the DdmD dimer (PDB: 9EZX), EMDB: EMD-50090) and the DdmD-DdmE holo-complex (PDB: 9EZY), EMDB: EMD-50091) have been deposited in the PDB and EMDB databases.

## Methods

### Bioinformatic analysis of pAgo orthologs

The prokaryotic orthologs used for the bioinformatic analysis were retrieved as described previously by Koopal et al. (*30*). In addition, DdmE was used as an input for the basic local alignment search tool of NCBI with a 60% sequence identity cutoff (*31*). All sequences were combined and aligned using MAFFT (v7.490) (*32*), after which a phylogenetic tree was generated using IQ-tree (v2.1.3) (*33*) with automated model selection (*34*) and visualized using FigTree (version 1.4.4). Structural models of different pAgo orthologs were generated using the Alphafold algorithm with the Colabfold environment (*35, 36*).

### Plasmid DNA constructs and site-specific mutants

For heterologous expression, the DNA sequences of *Vibrio cholerae* DdmD (Uniprot: Q9KR72) and DdmE (Uniprot: Q9KR73) were inserted into the 1B (Addgene: 29653) and 2HR-T (Addgene: 29718) plasmids using ligation-independent cloning (LIC), resulting in a construct that carries an N-terminal hexahistidine tag followed by a tobacco etch virus (TEV) protease cleavage site or a construct that carries an N-terminal hexahistidine-twin-strep-tactin tag followed by a TEV protease cleavage site, respectively. Site specific mutations were introduced by QuickChange mutagenesis or by inverse PCR. Plasmids were purified using the GeneJET plasmid miniprep kit (Thermo Fisher Scientific) and insertion and mutagenesis were verified by Sanger sequencing.

### Expression and purification of DdmE and StrepII-DdmE

Hexahistidine-tagged and Strep-tactin-tagged DdmE proteins were expressed in *E. coli* BL21-star cells. Cultures were grown at 37 °C, 130 rpm until they reached an OD_600_ of 0.6, after which the cultures were incubated on ice for 1 hour. Next, protein expression was induced with 0.25 mM isopropyl-β-D-thiogalactopyranoside (IPTG) and continued for 16 h at 18 °C, 130 rpm. Cells were harvested by centrifugation and resuspended in buffer A (20 mM Tris-HCl pH 8.0, 500 mM NaCl, 5 mM imidazole, 1 μg mL^−1^ pepstatin, 200 μg mL^−1^ AEBSF), followed by lysis in a Maximator cell homogenizer at 1,500 bar and 4 °C. The lysate was cleared by centrifugation at 10,000g for 30 min at 4 °C and applied to 15-mL equilibrated Ni-NTA beads (Qiagen). The Ni-NTA column was washed with 150 mL of buffer B (20 mM Tris-HCl pH 8.0, 500 mM NaCl, 10 mM imidazole), followed by 30 mL of buffer C (20 mM Tris-HCl pH 8.0, 150 mM NaCl, 10 mM imidazole). Proteins were eluted with five fractions of 15 mL of buffer D (20 mM Tris-HCl pH 8.0, 150 mM NaCl, 250 mM imidazole). Protein containing elution fractions were pooled and loaded onto two equilibrated 5-mL HiTrap Heparin HP columns (Cytiva) coupled in tandem and eluted with a linear gradient of buffer E (20 mM Tris-HCl pH 8.0, 1 M NaCl). Protein containing elution fractions were pooled and dialyzed overnight against buffer F (20 mM Tris-HCl pH 8.0, 250 mM NaCl, 1 mM DTT) in the presence of TEV protease. For proteins containing a Strep-tactin tag TEV protease was omitted from the dialysis. To remove uncleaved DdmE proteins, dialized proteins were supplemented with 10 mM Imidazole and ran over 7.5-mL equilibrated Ni-NTA beads (Qiagen). The flowthrough was collected and concentrated using 30 kDa molecular weight cut-off centrifugal filters (Merck Millipore) and further purified by size-exclusion chromatography using a Superdex 200 (16/600) column (Cytiva) equilibrated in buffer F. Purified proteins were concentrated to 4 mg mL^−1^, flash frozen in liquid nitrogen and stored at −80 °C until further use.

### Expression and purification of DdmD

Hexahistidine-tagged DdmD proteins were expressed in *E. coli* BL21-star cells. Cultures were grown at 37 °C, 130 rpm until they reached an OD_600_ of 0.6, after which the cultures were incubated on ice for 1 hour. Next, protein expression was induced with 0.25 mM isopropyl-β-D-thiogalactopyranoside (IPTG) and continued for 16 h at 18 °C, 130 rpm. Cells were harvested by centrifugation and resuspended in buffer A (20 mM Tris-HCl pH 8.0, 500 mM NaCl, 5 mM imidazole, 1 μg mL^−1^ pepstatin, 200 μg mL^−1^ AEBSF), followed by lysis in a Maximator cell homogenizer at 1,500 bar and 4 °C. The lysate was cleared by centrifugation at 10,000g for 30 min at 4 °C and applied to 15-mL equilibrated Ni-NTA beads (Qiagen). The Ni-NTA column was washed with 150 mL of buffer B (20 mM Tris-HCl pH 8.0, 500 mM NaCl, 10 mM imidazole). Proteins were eluted with five fractions of 15 mL of buffer C (20 mM Tris-HCl pH 8.0, 500 mM NaCl, 250 mM imidazole). Protein containing elution fractions were pooled and dialyzed overnight against buffer D (20 mM Tris-HCl pH 8.0, 350 mM NaCl, 1 mM DTT) in the presence of TEV protease. To remove uncleaved DdmD proteins, dialized proteins were supplemented with 10 mM Imidazole and ran over 7.5-mL equilibrated Ni-NTA beads (Qiagen). The flowthrough was collected and and concentrated using 100 kDa molecular weight cut-off centrifugal filters (Merck Millipore) and further purified by size-exclusion chromatography using a Superdex 200 (16/600) column (Cytiva) equilibrated in buffer D. Purified proteins were concentrated to 7 mg mL^−1^, flash frozen in liquid nitrogen and stored at −80 °C until further use.

### In vitro co-precipitation assays

Prior to grid preparation for cryo-EM, DdmE was loaded with a guide by mixing 10 μm of StrepII-DdmE with 13 μm DNA guide in a buffer containing 20 mM Tris-HCl pH 8.0, 250 mM NaCl, 5 mM MgCl_2_ and 0.05% Tween20. After 15 minutes of incubation at 37°C, 13 μm DNA target was added to the sample and incubated for 45 minutes at 37°C. Next, 50 μL of equilibrated Strep-tactin beads (IBA-life sciences) were added to the sample and incubated for 30 minutes at 4°C on a rotating wheel. Unbound proteins and DNA were removed by washing the beads were 3 times with a buffer containing 20 mM Tris-HCl pH 8.0, 250 mM NaCl, and 0.05% Tween20. Subsequently, 15 μm of DdmD was added to the beads and incubated for 30 minutes at room temperature on a rotating wheel. Unbound DdmD proteins were removed by washing the beads four times with a buffer containing 20 mM Tris-HCl pH 8.0, 250 mM NaCl and DdmDE complexes were eluted by incubating the beads for 5 minutes at 4°C with elution buffer containing 20 mM Tris-HCl pH 8.0, 250 mM NaCl and 2.5 mM Desthiobiotin.

### Electrophoretic mobility shift assays

For fluorescence polarization binding assays, DdmE was loaded with a guide by mixing 2.5 μm DdmE with 2.5 μm of DNA guide (Integrated DNA technologies, IDT) in a buffer containing 20 mM Tris-HCl, 250 mM NaCl, 5 mM MgCl 0.05% Tween20 and incubating the sample for 15 minutes at at 37°C. Next, 25 nM of fluorescently labelled single-stranded DNA substrate (IDT) was mixed with 2 μM unlabeled competitor DNA (IDT) and 1 μM of guide loaded DdmE in a binding buffer containing 20 mM Tris-HCl, 150 mM NaCl, 0.05% Tween20 and incubated for 30 minutes at 37°C. After incubation, samples were resolved using a 6% Native PAGE gel and scanned with the Typhoon trio (GE healthcare).

### *In vitro* plasmid degradation assays

For in vitro plasmid degradation assays, DdmE was loaded with a 14-nt phosphorylated DNA guide by mixing 10 μm DdmE with 15 μm of DNA guide (IDT) in a buffer containing 20 mM Tris-HCl, 100 mM NaCl, 5 mM MgCl_2_, and 5% glycerol and incubating the sample for 30 minutes at 37°C. Next, 10 nM of gel purified pUC19 plasmid was mixed with 1 μM of guide loaded DdmE in a binding buffer containing 20 mM Tris-HCl, 150 mM NaCl, 5% glycerol, 5% PEG8000 and incubated 18 hours at 20°C. After incubation, 2 μM of DdmD, 5 mM MgCl_2_, 5 mM MnCl_2_ and 0.5 mM ATP were added to the sample and incubated for the indicated times at 37°C, after which the samples were quenched by the addition of 0.5M EDTA and 1 μL Proteinase K and incubated for 10 minutes at 50°C. Next, samples were mixed with 6x DNA loading dye (10 mM Tris-HCl (pH 7.6), 60 mM EDTA, 60% Glycerol, 0.03% Bromophenol blue and 0.03% Xylene cyanol FF) and loaded onto a 0.8% agarose gel. Gels were run for 90 minutes at 100 V, followed by imaging with an ChemiDoc Imaging System (Biorad).

### General methods for *in vivo* assays

All *V. cholerae* strains are derivatives of the 7^th^ pandemic O1 El Tor (Inaba) strain A1552 (*37*). *E. coli* strain S17-1 λ*pir* was used for the propagation of plasmids with the conditional R6K origin of replication and for bacterial mating. Bacteria were cultured in Lysogeny Broth (LB-Miller; 10g/l NaCl, Carl Roth, Switzerland) at either 30°C (*V. cholerae*) or 37°C (*E. coli*), and where needed, antibiotic selection was applied using Ampicillin (100 μg/ mL) or Kanamycin (75 μg/ mL).

### V. cholerae strain construction

Scar-less and marker-less genetic manipulations of *V. cholerae* were performed by allelic exchange using the counter-selectable plasmid pGP704-Sac28, as previously described (*38*). In brief, following bi-parental mating with an *E. coli* donor strain, *V. cholerae* was selected using thiosulfate citrate bile salts sucrose (TCBS; Sigma-Aldrich) agar supplemented with Ampicillin (100 μg/ mL). Subsequently, a SacB-mediated counter-selection was performed on NaCl-free LB media containing 10% sucrose and colonies were screened for successful exchanges. Site-directed mutants of plasmid pGP704-TnAraC-*ddmDE* were obtained by inverse PCR. All genetic constructs were validated by PCR and Sanger or full plasmid Nanopore sequencing (Microsynth AG, Switzerland). Plasmids were introduced into *V. cholerae* by conjugation or electroporation and into chemically competent cells of *E. coli*, as previously described (*13*).

### *V. cholerae* plasmid elimination assay

Plasmid stability assays were performed as previously described (*13*). In brief, plasmid stability in *V. cholerae* was determined by measuring the retention of plasmid pSa5Y-Kan over ∼50 generations without selection pressure. Cells were grown at 30°C under shaking conditions (180rpm).

### *E. coli* plasmid-based plasmid elimination assay

For *E. coli-*based plasmid stability assays, the DdmDE activity was evaluated in strain S17-1λ*pir*. In this assay, the *ddmD* and *ddmE* genes were encoded by plasmid pGP704-TnAraC-*ddmDE*, which carries an arabinose-inducible promoter (*P*_BAD_) and becomes self-eliminating upon arabinose-induced expression of *ddmDE*. Strains carrying either pGP704-TnAraC (negative control), pGP704-TnAraC-*ddmDE* (positive control) or pGP704-TnAraC-*ddmDE* derivatives encoding variants in either DdmD or DdmE, were cultured overnight with selection (Ampicillin, 100 μg/ mL) and diluted to an O.D._600_ of 0.0025 in fresh LB, and grown without selection pressure for ∼10 generation at 37°C, 180 rpm, in the absence and presence 0.2% arabinose. Subsequently, cells were serially diluted in phosphate buffered saline (PBS), and 5 μL of each dilution spotted on LB agar plates in the absence and presence of selection (Amp 100 μg/ml). Plates were incubated over night at 37°C and plasmid stability was measured by comparing the ratio of the antibiotic resistant (*i*.*e*. plasmid-carrying) colony forming units against the total number of bacteria.

### SEC-MALS analysis of DdmD and DdmE

For SEC-MALS purified proteins were thawed and centrifuged at 21,000g for 10 min at 4 °C, after which the supernatant was filtered with a 0.1 μm centrifugal filter (Merck Millipore). The sample was injected at a concentration 1 mg mL^-1^ onto a Superdex 200 (3.2/300) column (Cytiva) that was equilibrated in a buffer containing 20 mM Tris-HCl pH 8.0, 250 mM NaCl, 1 mM DTT. After SEC, the sample passed a miniDAWN TREOS (599-TS) multiple angle light scattering detector (Wyatt Technology) and a Optilab rEX (329-rEX) refractive index detector (Wyatt Technology). The light source of the RI detector was a G1315B DAD UV detector (Agilent) and wavelength of the laser in the light scattering instrument was set at 658.9 nm. Prior to the sample a bovine serum albumin (BSA) sample was run as a calibration standard. Data collection and analysis were performed in the ASTRA 6.1 software (Wyatt Technology) with the refractive index of the solvent, the refractive index increment (∂n/∂c) and the viscosity defined as 1.331, 0.185 mL g^-1^ and 0.8945 cP, respectively.

### DdmD DNA unwinding assays

Prior to the cleavage assays, synthetic dsDNA targets (Integrated DNA technologies) were annealed in a buffer containing 10 mM Tris-HCl, 50 mM NaCl, using a thermocycler (Biorad) and stored at -20°C until further use. For DNA unwinding assays, 50 nM of DNA target was mixed with 500 nM of the DdmD in a buffer containing 20 mM Tris-HCl, 125 mM NaCl 5 mM MgCl_2_ and 0.05% Tween20 and incubated for 15 minutes at 4°C. After the incubation, samples were transferred to a 96 well plate and 0.5 mM was injected while measuring the fluorescence intensity using a Pherastar plate reader (BMG labtech). For endpoint measurements, samples were incubated with ATP for 5 minutes at room temperature pior to measuring the fluorescence intensity.

### Sample preparation and cryo-EM data collection of apo DdmD

Prior to grid preparation for cryo-EM, thawed protein samples were purified by size-exclusion chromatography using a Superdex 200 (16/600) column (GE Healthcare) in equilibrated in a buffer containing 20 mM Tris-HCl pH 8.0, 350 mM NaCl. Peak fractions were concentrated to 1.5 mg mL^-1^ using 100 kDa molecular weight cut-off centrifugal filters (Merck Millipore). To each 200-mesh holey carbon grid (Au R1.2/1.3, Quantifoil Micro Tools), 2.5 μL of sample was applied and blotted for 3 s at 80% humidity and 4 °C. Grids were plunge frozen in liquid ethane, using a Vitrobot Mark IV plunger, FEI) and stored in liquid nitrogen until cryo-EM data collection. Cryo-EM data collection was performed on a FEI Titan Krios microscope equipped with a Gatan K3 direct electron detector (University of Zurich) operated at 300 kV in super-resolution counting mode. Data acquisition was performed using the EPU Automated Data Acquisition Software for Single Particle Analysis from ThermoFisher with three shots per hole at defocus range of −1.0 μm to −2.4 μm (0.2-μm steps). The final dataset comprised a total of 9,031 micrographs at a calibrated magnification of 130,000x and a super-resolution pixel size of 0.325 Å. Micrographs were exposed for 1.01s with a total dose of 59.98 e^−^ Å^−2^ over 42 subframes.

### Data processing and model building of apo DdmD

Cryo-EM data was processed using cryoSPARC (v4.4.1) (*39*). The 9,031 micrographs were imported and motion-corrected with patch motion correction (multi) after which, the CTF values of the micrographs were estimated using patch CTF estimation (multi). Next, an initial set of particles was picked with blob picker using a circular blob and a minimum and maximum particle diameter of 50 and 150 Å, respectively. After extraction of the particles with a box size of 572x572 pixels, particles were subjected to 2D classification to generate templates for picking (5 templates).

After template-based picking with a particle diameter of 150 Å, particles were extracted and subjected to 2D classification with a circular mask of 160 Å. Classes with defined particles were selected, resulting in a total of 2,023,352 particles, which were used to generate two *ab initio* models of which one was used for heterogeneous refinement with five classes. Classes were inspected visually using UCSF Chimera(*40*), and the particles and volume of the best class were subjected to a second round of *ab initio* model generation with four classes and the class similarity setting set to 0.9. The particles and volume of the best class were used as an input for per-particle motion correction and subsequently refined using non-uniform refinement with optimization of CTF parameters and C2 symmetry enabled. The final map was sharpened with a B-factor of -85. The local resolution was estimated based on the resulting map using the local resolution function of cryoSPARC and plotted on the map using UCSF Chimera(*40*).

The structural model of Apo DdmD was built *de novo* in Coot (V0.9.2) (*41*) and was refined over multiple rounds using Phenix (*42, 43*). Real-space refinement was performed with the global minimization, atomic displacement parameter (ADP) refinement and secondary structure restrains enabled. The quality of the atomic model, including protein geometry, Ramachandran plots, clash analysis and model cross-validation, was assessed with MolProbity and the validation tools in Phenix (*42–45*). The refinement statistics of the final model are listed in Supplementary Table S1. Figures of maps, models and the calculations of map contour levels were generated using ChimeraX(*40*).

### Sample preparation and cryo-EM data collection of DdmDE complex

Prior to grid preparation for cryo-EM, DdmE was loaded with a guide by mixing 10 μm of StrepII-DdmE with 13 μm DNA guide in a buffer containing 20 mM Tris-HCl pH 8.0, 225 mM NaCl, 5 mM MgCl_2_ and 0.05% Tween20. After 15 minutes of incubation at 37°C, 13 μm DNA target was added to the sample and incubated for 45 minutes at 37°C. Next, 50 μL of equilibrated Strep-tactin beads (IBA-life sciences) were added to the sample and incubated for 30 minutes at 4°C on a rotating wheel. Unbound proteins and DNA were removed by washing the beads were 3 times with a buffer containing 20 mM Tris-HCl pH 8.0, 225 mM NaCl, and 0.05% Tween20. Subsequently, 15 μm of DdmD was added to the beads and incubated for 30 minutes at room temperature on a rotating wheel. Unbound DdmD proteins were removed by washing the beads four times with a buffer containing 20 mM Tris-HCl pH 8.0, 225 mM NaCl and DdmDE complexes were eluted by incubating the beads for 5 minutes at 4°C with elution buffer containing 20 mM Tris-HCl pH 8.0, 225 mM NaCl and 2.5 mM Desthiobiotin. Eluted proteins were concentrated to 2.5 mg mL^-1^ using 100 kDa molecular weight cut-off centrifugal filters (Merck Millipore). To each 200-mesh holey carbon grid (Au R1.2/1.3, Quantifoil Micro Tools), 2.5 μL of sample was applied and blotted for 3 s at 80% humidity and 4 °C. Grids were plunge frozen in liquid ethane, using a Vitrobot Mark IV plunger, FEI) and stored in liquid nitrogen until cryo-EM data collection. Cryo-EM data collection was performed on a FEI Titan Krios microscope equipped with a Gatan K3 direct electron detector (University of Zurich) operated at 300 kV in super-resolution counting mode. Data acquisition was performed using the EPU Automated Data Acquisition Software for Single Particle Analysis from ThermoFisher with three shots per hole at defocus range of −1.0 μm to −2.4 μm (0.2-μm steps). The final dataset comprised a total of 9,031 micrographs at a calibrated magnification of 130,000x and a super-resolution pixel size of 0.325 Å. Micrographs were exposed for 1.01s with a total dose of 60.99 e^−^ Å^−2^ over 42 subframes.

### Data processing and model building of DdmDE complex

Cryo-EM data was processed using cryoSPARC (v4.4.1) (*39*). The 9,974 micrographs were imported and motion-corrected with patch motion correction (multi) after which, the CTF values of the micrographs were estimated using patch CTF estimation (multi). Next, an initial set of particles was picked with blob picker using a circular blob and a minimum and maximum particle diameter of 50 and 150 Å, respectively. After extraction of the particles with a box size of 572x572 pixels, particles were subjected to 2D classification to generate templates for picking (5 templates).

After template-based picking with a particle diameter of 150 Å, particles were extracted and subjected to 2D classification with a circular mask of 180 Å. Classes with defined particles were selected, resulting in a total of 1,283,702 particles, which were used to generate two *ab initio* models of which one was used for heterogeneous refinement with four classes. Classes were inspected visually using UCSF Chimera(*40*), and the particles and volume of the best class were used as an input for per-particle motion correction and subsequently refined using non-uniform refinement with optimization of per-particle defocus and CTF parameters. The final map was sharpened with a B-factor of -65. The local resolution was estimated based on the resulting map using the local resolution function of cryoSPARC and plotted on the map using UCSF Chimera (*40*).

The structural model of the DdmDE was built *de novo* in Coot (V0.9.2) (*41*) and was refined over multiple rounds using Phenix (*42, 43*). Real-space refinement was performed with the global minimization, atomic displacement parameter (ADP) refinement and secondary structure restrains enabled. The quality of the atomic model, including protein geometry, Ramachandran plots, clash analysis and model cross-validation, was assessed with MolProbity and the validation tools in Phenix (*42–45*). The refinement statistics of the final model are listed in Supplementary Table S1. Figures of maps, models and the calculations of map contour levels were generated using ChimeraX (*40*).

