## Supplemental Figures for "Molecular mechanism of plasmid elimination by the DdmDE defense system"

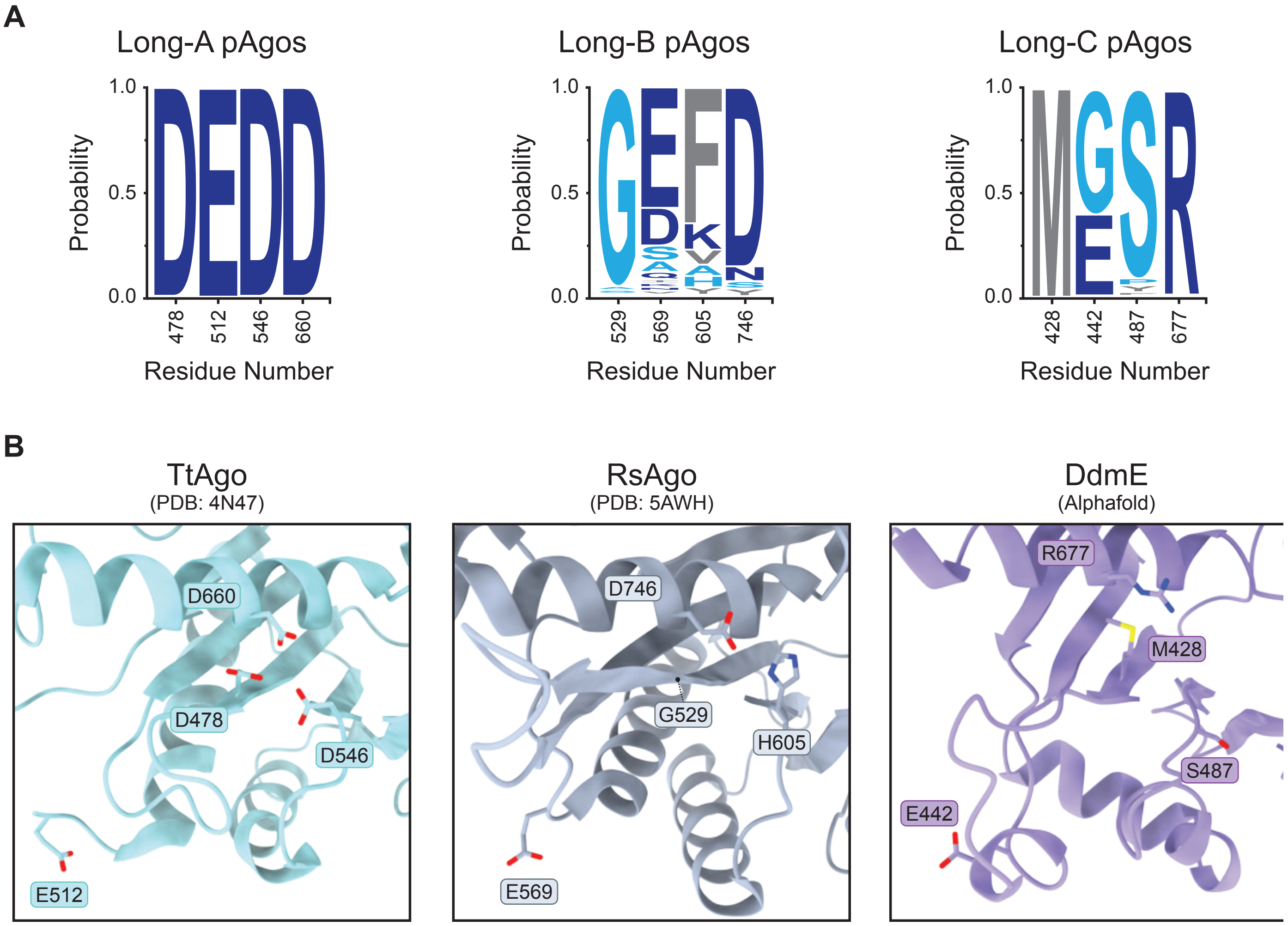


**Figure S1: Comparison of PIWI domains within long pAgos.**

**(A)** Sequence conservation analysis of active site residues in different clades of long pAgo proteins. Figure was generated using WebLogo (*46*). **(B)** Structural comparison of pAgos that represent the clades Long-A (*Thermus thermophilus* Argonaute, TtAgo, PDB: 4N47 (*47*)), Long-B (*Rhodobacter sphaeroides* Argonaute, RsAgo, PDB: 5AWH (*48*)) and Long-C *Vibrio cholerae* DdmE, AlphaFold model (*35*, *36*)).

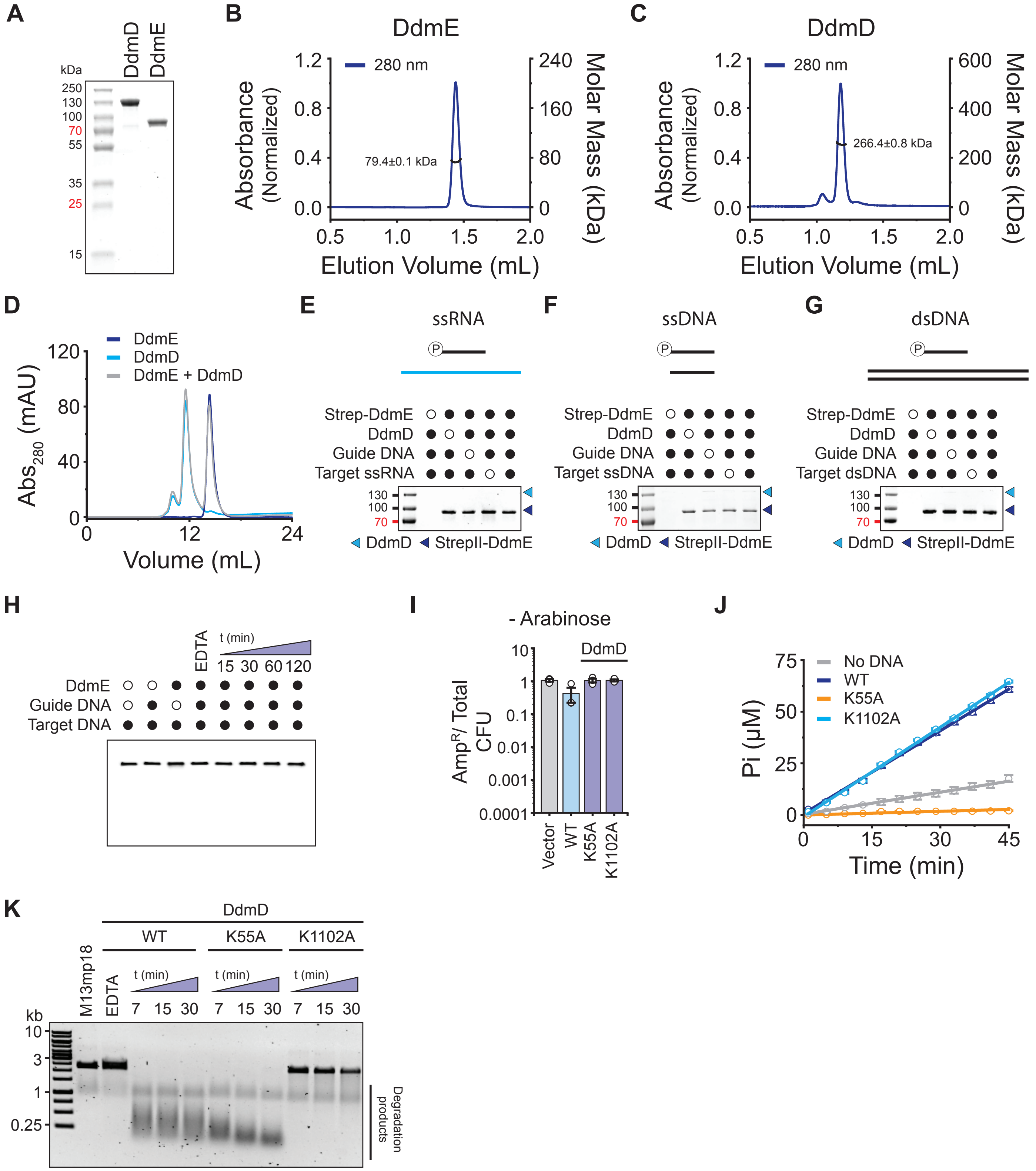


**Figure S2: Biochemical characterization of the DdmDE system.**

**(A)** Analysis of purified recombinantly-expressed DdmD and DdmE by SDS-PAGE. **(B)** Size-exclusion chromatography coupled to multi-angle static light scattering (SEC-MALS) analysis of DdmE. **(C)** SEC-MALS analysis of DdmD. **(D)** Size-exclusion chromatography analysis of DdmD, DdmE and their mixture. **(E)** *In vitro* pull-down experiment with purified DdmD and StrepII-DdmE proteins in the presence of 14-nt 5'-phosphorylated DNA guide and an complementary ssRNA target. Strep-tactin beads were washed to remove unbound DdmD, and bound proteins were analyzed by SDS-PAGE and stained with Coomassie blue. **(F)** *In vitro* pull-down experiment with purified DdmD and StrepII-DdmE proteins in the presence of 14-nt 5'-phosphorylated DNA guide and a complementary 22-nt ssDNA target. **(G)** *In vitro* pull-down experiment with purified DdmD and StrepII-DdmE proteins in the presence of 14-nt 5'-phosphorylated DNA guide and a 50-bp perfectly complementary dsDNA target. **(H)** *In vitro* nuclease activity assays using fluorescently labeled ssDNA oligonucleotide complementary to the guide DNA. Samples were resolved by denaturing PAGE. **(I)** *In vivo* plasmid elimination assay in the absence of the inducer arabinose. *E. coli* strains harbor vectors encoding WT DdmDE, WT DdmE and DdmD mutants or an empty vector control. Data represents mean fraction of plasmid-bearing colony forming units (CFU) ± SEM of three independent replicates (n=3). **(J)** ATPase activity of DdmD variants on ssDNA. **(K)** *In vitro* plasmid degradation assay using a single-stranded plasmid substrate in the presence of WT DdmD or DdmD proteins carrying inactivating mutations in the helicase (K55A) or nuclease (K1102A) domains. Negative controls were incubated for the entire duration of the experiment.

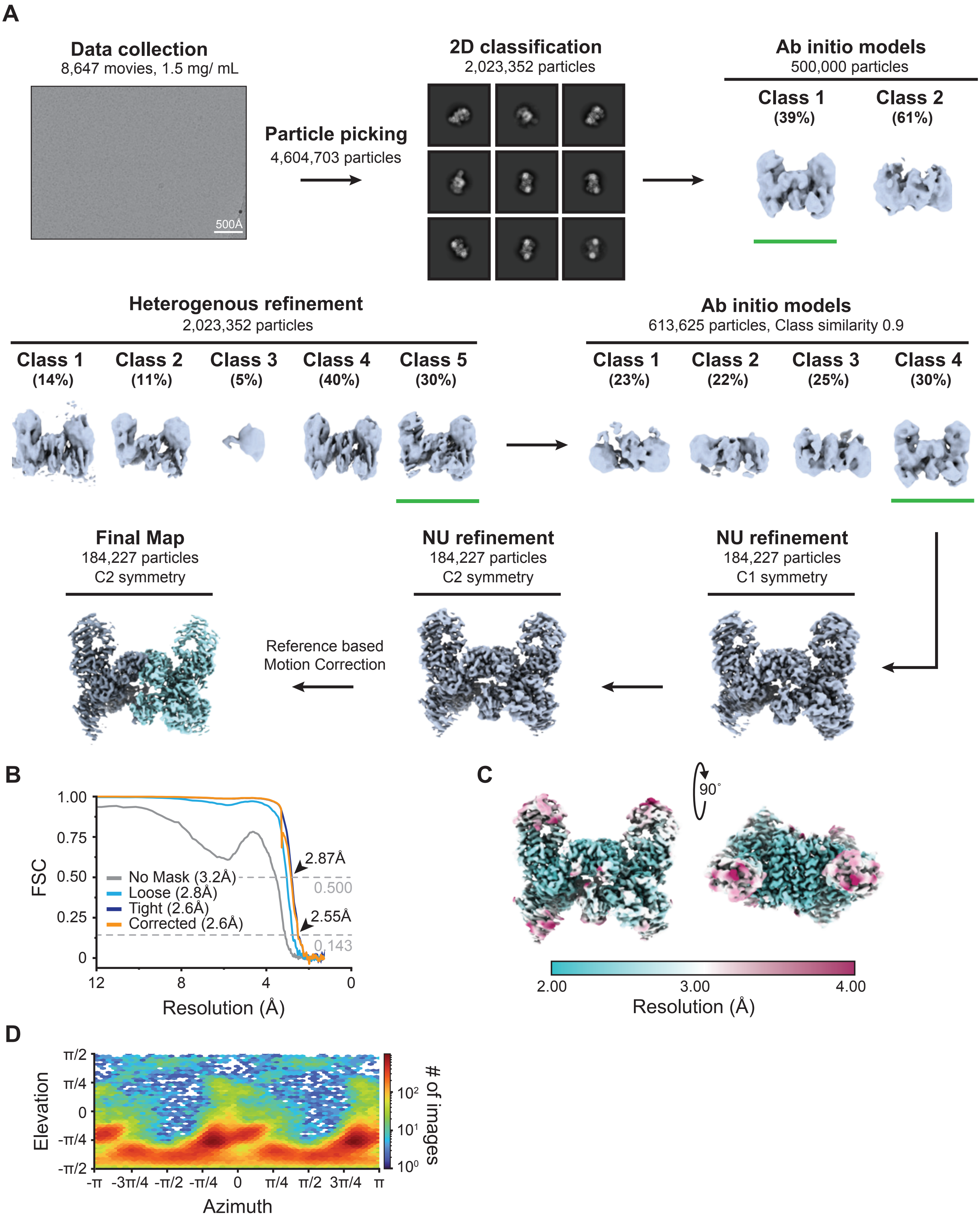


**Figure S3: Cryo-EM processing workflow for the DdmD dimer.**

**(A)** Cryo-EM data collection and processing workflow for the DdmD dimer. **(B)** Fourier Shell Correlation (FCS) determined from two independently refined half-maps. The gold standard cut-off (FCS=0.143) is marked with an arrow. **(C)** Local resolution estimation on the final cryo-EM density map of the DdmD dimer. **(D)** Euler diagram showing orientation distribution of final cryo-EM reconstruction.

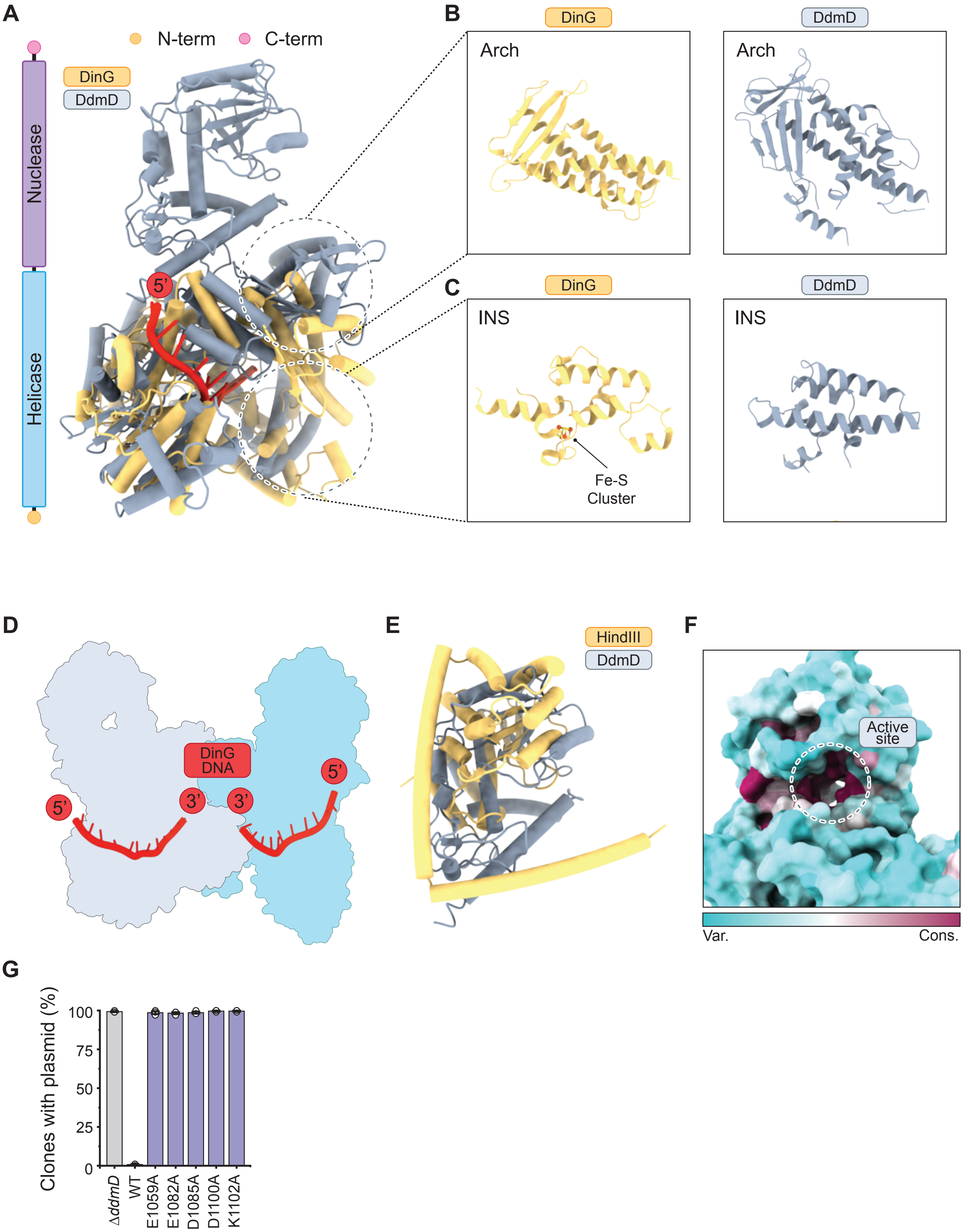


**Figure S4: Molecular details of the DdmD monomer.**

**(A)** Structural superposition of DdmD (light grey) with DinG bound to single-stranded DNA (DinG orange, DNA red, PDB: 6FWS) (*16*). **(B)** Close-up view of the DinG (orange, PDB: 6FWS) and DdmD (light grey) arch domains. **(C)** Close-up view of the DinG (orange, PDB: 6FWS) and DdmD (light grey) INS domains. **(D)** Schematic diagram of the DdmD dimer with bound DNA, superimposed with ssDNA as bound in the structure of DNA-bound DinG. The DdmD dimer is displayed as a surface outline, the DNA is displayed as cartoon (red). DinG protein has been omitted from the schematic for clarity. **(E)** Structural superposition of the C-terminal domain of DdmD (light grey) with nuclease domain of HindIII (orange, PDB: 3WVG) (*49*). **(F)** Side view of an DdmD monomer colored by sequence conservation using ChimeraX (*50*). The nuclease active site is indicated by a dotted circle. **(G)** *In vivo* plasmid elimination assay in *V. cholerae* strains expressing WT DdmDE, WT DdmE and DdmD mutants or a Δ*ddmD* control. Data represents the mean fraction of plasmid-bearing clones ± SEM of three independent

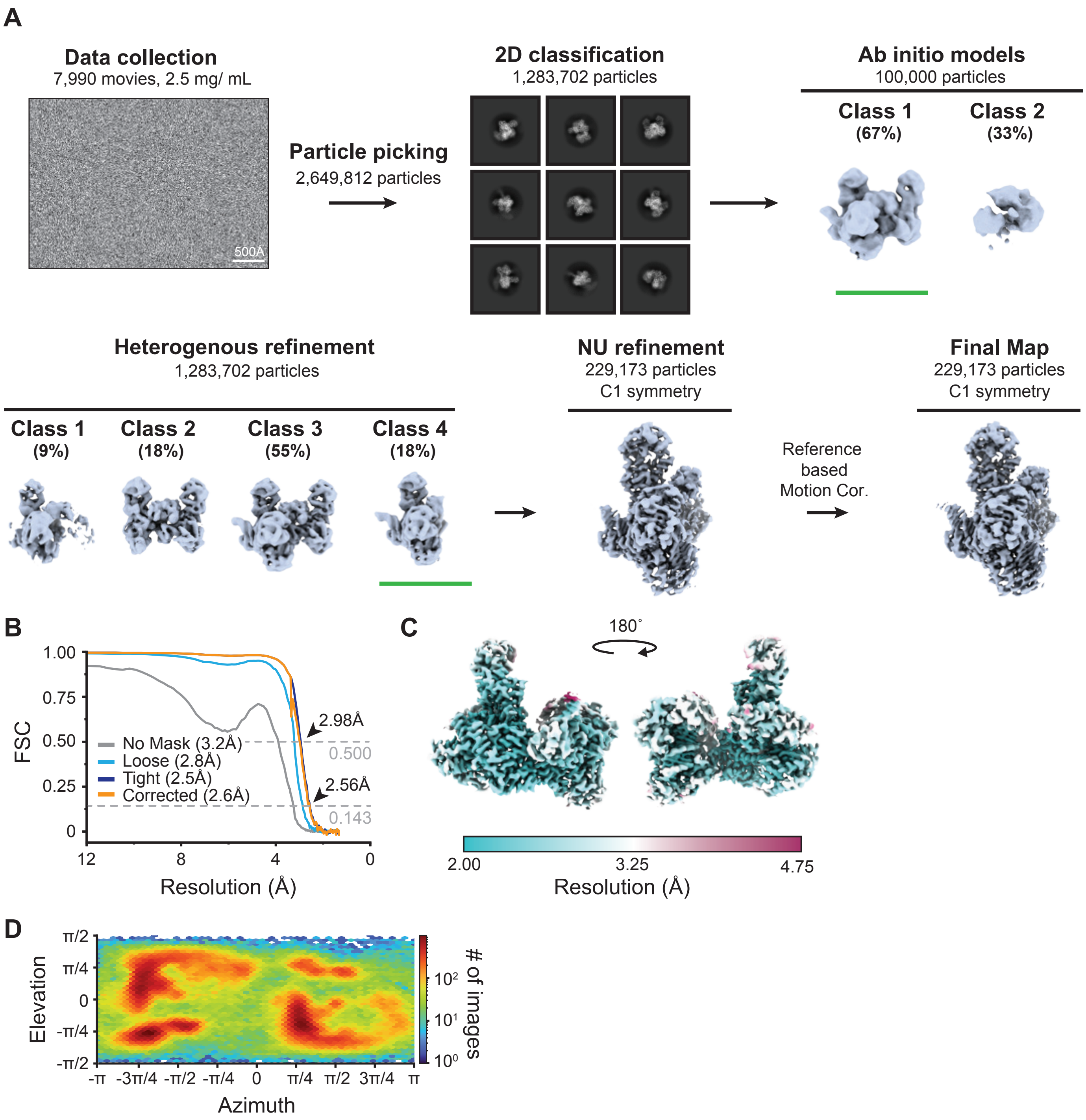


**Figure S5: Cryo-EM processing workflow for the DdmD-DdmE holocomplex.**

**(A)** Cryo-EM data collection and processing workflow for the DdmD-DdmE holocomplex. **(B)** Fourier Shell Correlation (FCS) determined from two independently refined half-maps. The gold standard cut-off (FCS=0.143) is marked with an arrow. **(C)** Local resolution estimation on the final cryo-EM density map of the DdmDE holo complex. **(D)** Euler diagram showing orientation distribution of final cryo-EM reconstruction.

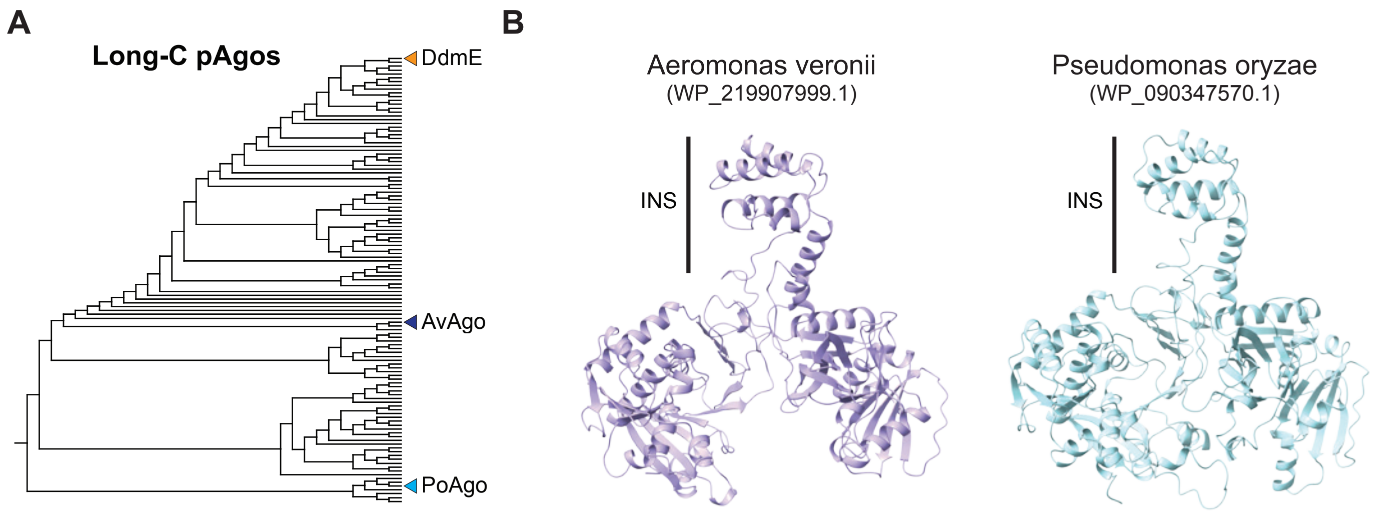


**Figure S6: Bioinformatic analysis Long-C pAgos.**

**(A)** Phylogenetic tree of Long-C pAgos. **(B)** AlphaFold structure predictions showing the INS domain is conserved across the Long-C pAgo family.

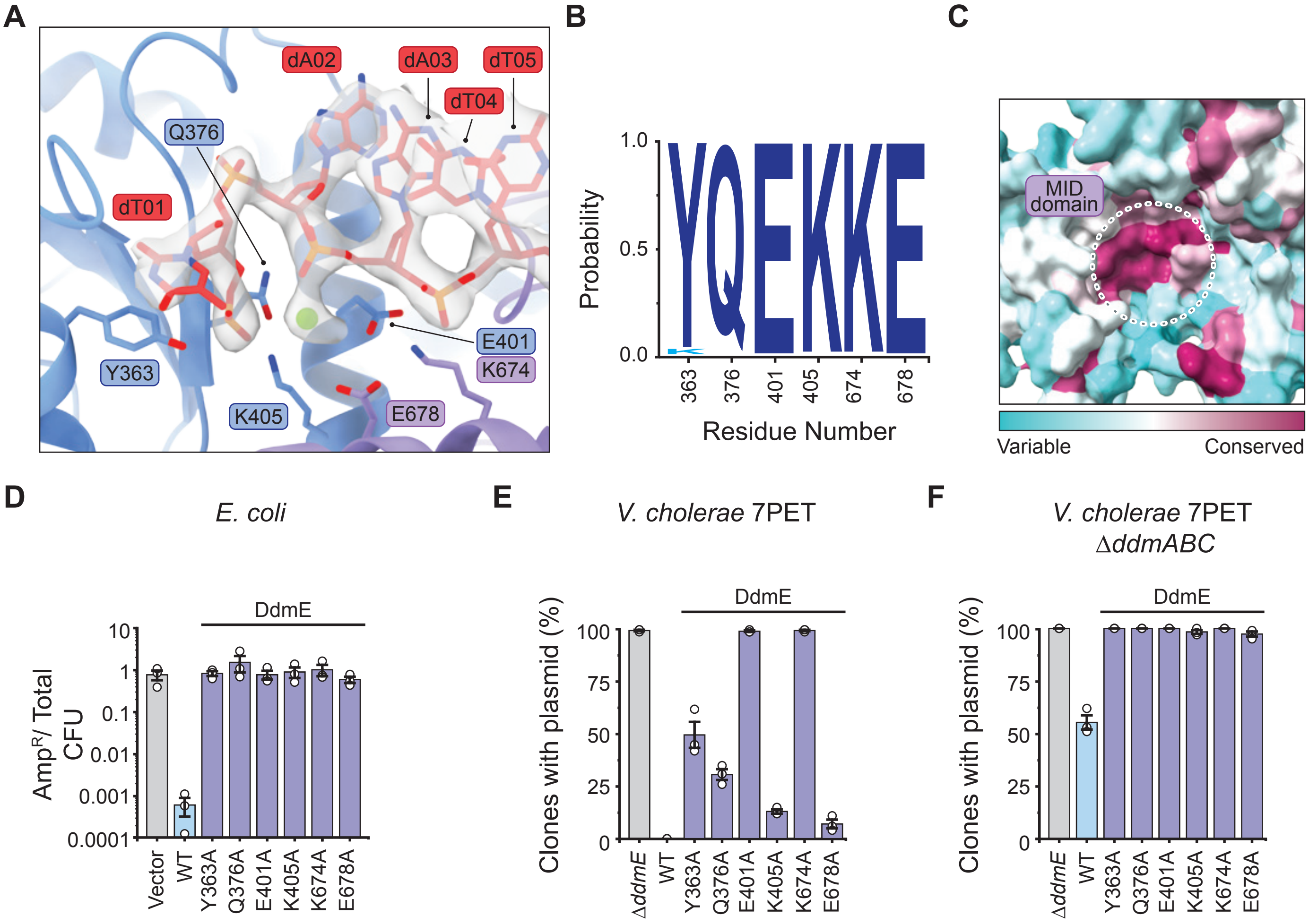


**Figure S7: Structural details of guide DNA binding by DdmE.**

(**A**) Close-up view of DdmE MID domain housing the 5' end of the guide DNA. Cryo-EM density for the DNA is shown in grey. (**B**) Sequence conservation analysis of key MID domain residues involved in guide DNA binding. Figure was generated using WebLogo (*46*). **(C)** Surface view of the MID domain colored by sequence conservation using ChimeraX (*50*). (**D**) *In vivo* plasmid elimination assay in *E. coli* strains expressing WT DdmDE, WT DdmD and DdmE MID domain mutants, or a vector-only control. Data represents mean plasmid-bearing colony forming units (CFU) ± SEM of three independent replicates (n=3). Experimental data was acquired simultaneously with data displayed in Fig 2F and share the same WT and vector-only controls. (**E**) *In vivo* plasmid elimination assay in 7PET *V. cholerae* strains expressing WT DdmDE, DdmE MID domain mutants, or a Δ*ddmE* control. Data represents the mean fraction of plasmid bearing clones ± SEM of three independent replicates (n=3). (**F**) *In vivo* plasmid elimination assay in Δ*ddmABC* mutant 7PET *V. cholerae* strains expressing WT DdmDE, DdmE MID domain mutants or a Δ*ddmE* control. Data represents the mean fraction of plasmid-bearing clones ± SEM of three independent replicates (n=3).

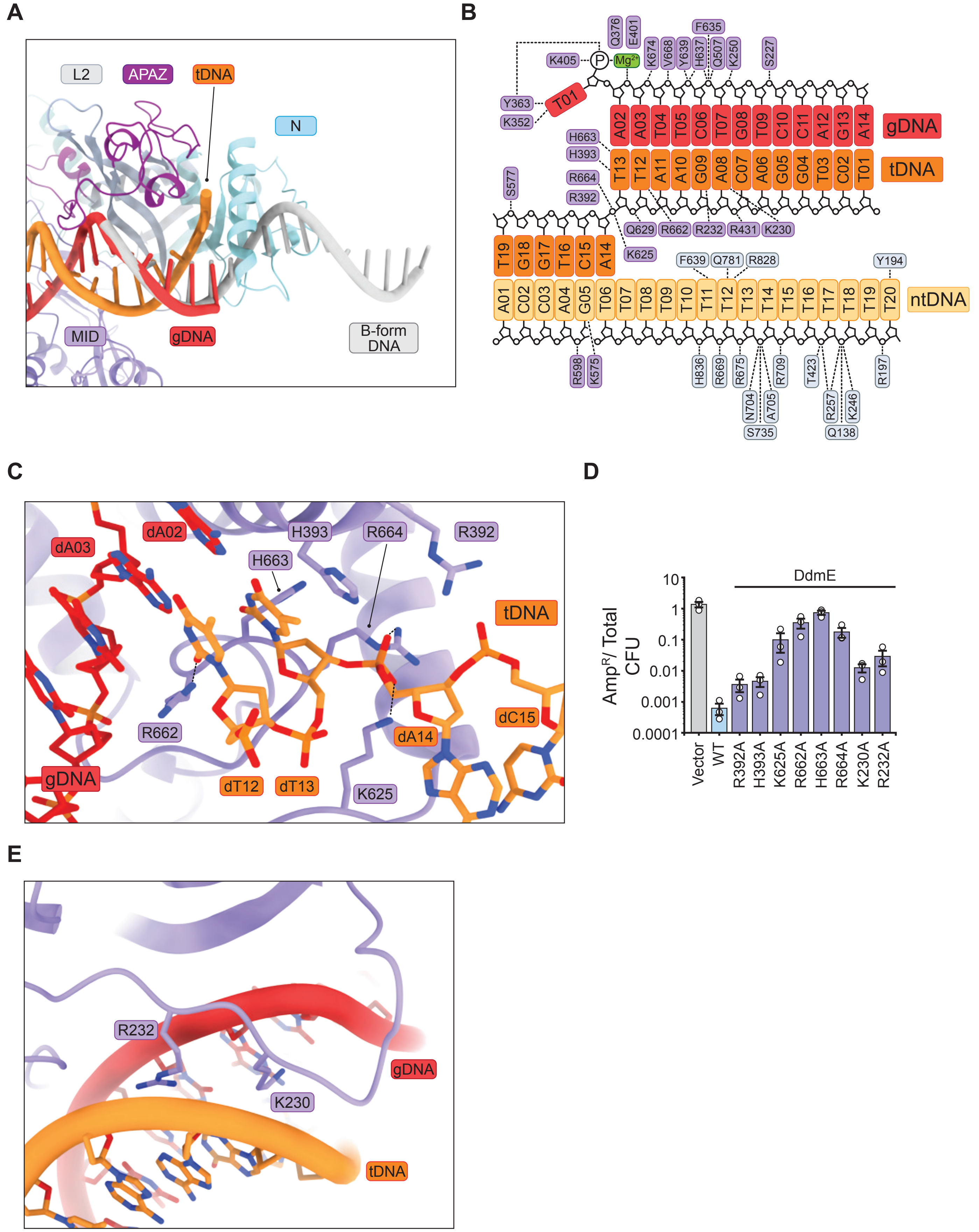


**Figure S8: Structural details of target DNA binding by DdmE**

**(A)** Structural superposition of the guide DNA (red) and ideal B-form single-stranded DNA (grey), indicating steric clash between DdmE and 3' terminal extension of the guide. **(B)** Schematic showing interactions of DdmD (grey) and DdmE (purple) the guide and target DNAs. **(C)** Close-up view of DdmE interactions at the proximal end of the dsDNA target (orange). Nucleotides 9-11 of the target DNA have been removed from the model for clarity. (**D**) *In vivo* plasmid elimination assay in *E. coli* strains expressing WT DdmDE, WT DdmD and DdmE mutants, or a vector-only control. Data represents mean fraction of plasmid-bearing colony forming units (CFU) ± SEM of three independent replicates (n=3). **(E)** Close-up view of the minor groove interrogation by the DdmE C-APAZ domain.

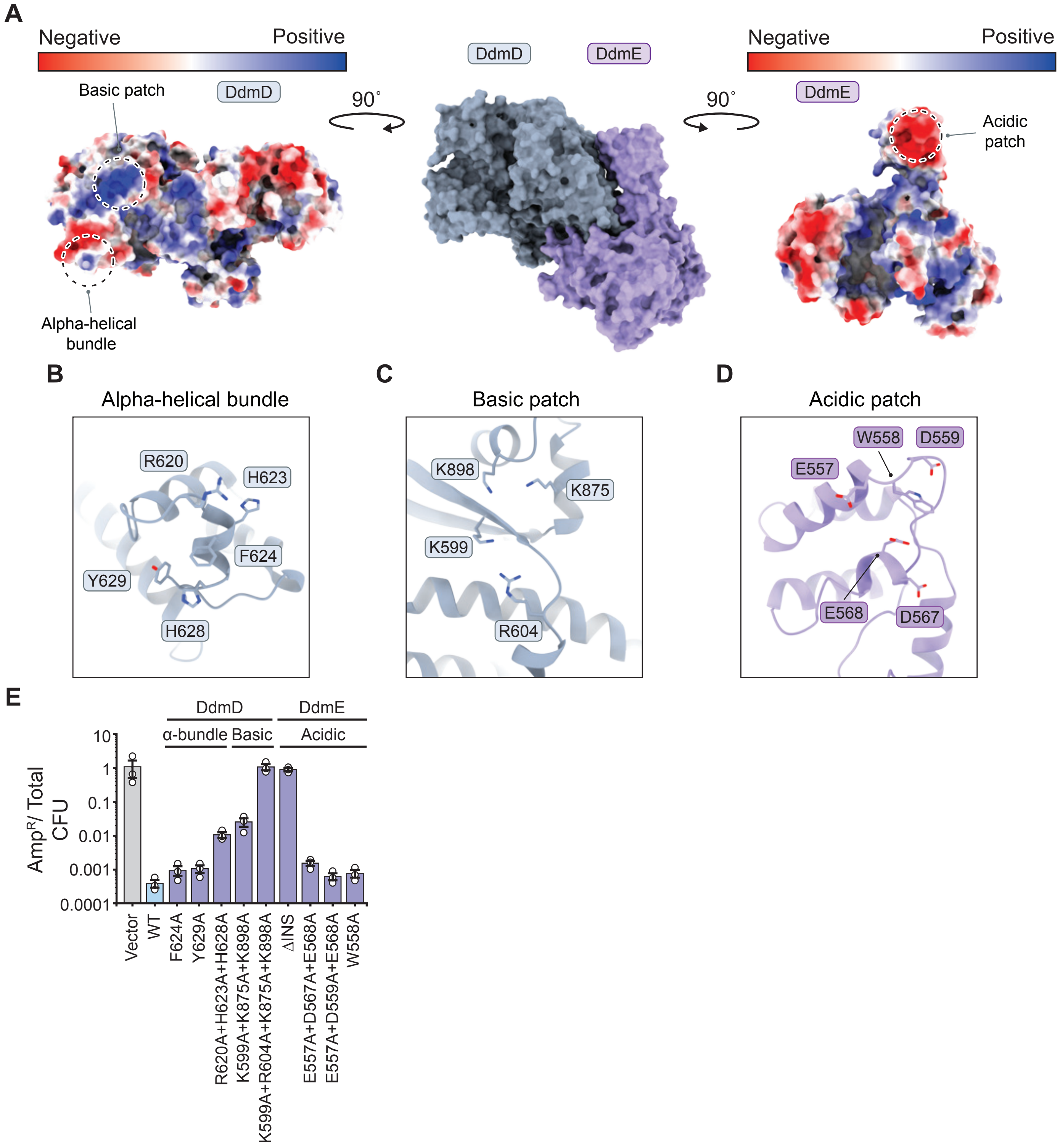


**Figure S9: Molecular details of the DdmD-DdmE interaction surfaces.**

**(A)** Orthogonal views of the DdmD-DdmE holocomplex. Middle: Surface view of the DdmD-DdmE complex. Right: Surface view of DdmD, colored by electrostatic potential. Left: Surface view of DdmE, colored by electrostatic potential (left). **(B)** Close-up view of the DdmD alpha-helical bundle. **(C)** Close-up view of basic patch in DdmD. **(D)** Close-up view of the acidic patch in the INS domain of DdmE. **(E)** *In vivo* plasmid elimination assay in *E. coli* strains expressing WT DdmDE, WT DdmE and DdmD mutants, DdmD mutants and WT DdmE, or a vector-only control. Data represents mean fraction of plasmid-bearing colony forming units (CFU) ± SEM of three independent replicates (n=3). Experimental data was acquired simultaneously with data displayed in Fig 4D and share the same WT and vector-only controls.

**Table S1: Cryo-EM data collection, refinement, and validation statistics for *Vibrio cholerae* DdmD and DdmD-DdmE complex**

| | *Vibrio cholerae* DdmD apo complex (EMDB: EMD-50090) (PDB: 9EZX) | *Vibrio cholerae* DdmD-DdmE holo complex (EMDB: EMD-50091) (PDB 9EZY) |
|---|---|---|
| **Data collection and processing** | | |
| Magnification | 130,000 x | 130,000 x |
| Voltage (kV) | 300 | 300 |
| Electron exposure (e–/Å$^2$) | 66.53 | 60.01 |
| Defocus range (μm) | -1.0 to -2.4 (-0.2 steps) | -1.0 to -2.4 (-0.2 steps) |
| Pixel size (Å) | 0.65 | 0.65 |
| Symmetry imposed | C2 | C1 |
| Initial particle images (no.) | 2,023,352 | 1,283,702 |
| Final particle images (no.) | 184,227 | 229,173 |
| Map resolution (Å) | 2.55 | 2.56 |
| FSC threshold | 0.143 | 0.143 |
| Map resolution range (Å) | 2.0 – 4.8 | 2.0 – 4.0 |
| **Refinement** | | |
| Model resolution (Å) | 2.5 | 2.6 |
| FSC threshold | 0.143 | 0.143 |
| Model resolution range (Å) | 2.5-2.7 | 2.5-2.9 |
| Map sharpening *B* factor (Å$^2$) | -85 | -65 |
| Model composition | | |
| Non-hydrogen atoms | 18585 | 15541 |
| Protein residues | 2302 | 1786 |
| Nucleotides | 0 | 52 |
| Ligands | 0 | MG:1 |
| *B* factors (Å$^2$) | | |
| Protein | 32.00/254.91/116.38 | 56.27/246.74/120.58 |
| Nucleotides | - | 24.84/170.31/75.21 |
| Ligand | - | 65.79/65.79/65.79 |
| R.m.s. deviations | | |
| Bond lengths (Å) | 0.005 | 0.003 |
| Bond angles (°) | 0.741 | 0.600 |
| Validation | | |
| MolProbity score | 1.32 | 1.29 |
| Clashscore | 4.71 | 4.59 |
| Poor rotamers (%) | 1.03 | 0.63 |
| Ramachandran plot | | |
| Favored (%) | 97.67 | 97.73 |
| Allowed (%) | 2.33 | 2.27 |
| Disallowed (%) | 0.00 | 0.00 |
